# A non-acidic method using hydroxyapatite and phosphohistidine monoclonal antibodies allows enrichment of phosphopeptides containing non-conventional phosphorylations for mass spectrometry analysis

**DOI:** 10.1101/691352

**Authors:** K. Adam, S. Fuhs, J. Meisenhelder, A. Aslanian, J. Diedrich, J. Moresco, J. La Clair, J.R. Yates, T. Hunter

## Abstract

Four types of phosphate-protein linkage generate nine different phosphoresidues in living organisms. Histidine phosphorylation is a long-time established but largely unexplored post-translational modification, mainly because of the acid-lability of the phosphoramidate bonds. This lability means that standard phosphoproteomic methods used for conventional phosphate esters (phospho-Ser/Thr/Tyr) must be modified to analyze proteins containing the phosphoramidate-amino acids - phospho-His/Arg/Lys. We show that a non-acidic method allows enrichment of non-conventional phosphoresidue-containing peptides from tryptic digests of human cell lines, using hydroxyapatite binding and/or immobilized 1-pHis and 3-pHis monoclonal antibodies for enrichment. 425 unique non-conventional phosphorylation sites (i.e. pHis, pLys and pArg) were detected with a high probability of localization by LC-MS/MS analysis and identified using a customized MaxQuant configuration, contributing to a new era of study in post-translational modification and cell signaling in humans. This is the first fully non-acidic method for phosphopeptide enrichment which uses immunoaffinity purification and remains compatible with mass spectrometry analysis for a wider coverage of potential protein phosphorylation events.

## Introduction

Phosphoproteomics has contributed to tremendous progress in the field of life science and revealed the extensive use of phosphorylation as one of the most important post-translational modifications in eukaryotic organisms. The identification and characterization of phosphorylation sites has been revolutionized by mass spectrometry (MS) with faster and more sensitive instruments. Progress in phosphosite-mapping with tandem mass spectrometry (MS/MS) has allowed improved localization and phosphorylation assignment to a single residue with high confidence. The development of bioinformatics tools for large-scale phosphoproteome data analysis was an equally important step and permitted the integration of different parameters searching for complex combinatorial possibilities ^1,2^.

Up until now, only one class of phosphate-bond containing phosphoamino acid, corresponding to the conventional serine, threonine and tyrosine phosphorylation (phosphate-ester or O-phosphate) has been systematically considered for phosphorylation site searching. But nine out of the 20 amino acids have the potential to be phosphorylated within the SONAtes covering the 4 categories of phosphate bonds: *S*-phosphate (thiophosphate), *O*-phosphate (phosphate-ester), *N*-phosphate (phosphoramidate) and *A*-phosphate (acyl-phosphate group). Phosphoramidate includes the histidine phosphorylation, well-defined in bacteria ^3,4^. Regardless of the nature of their chemical linkages, the presence of the phosphate group on all nine amino acids causes an increased mass of 79.97 Da, which can be detected by MS. During phosphopeptide fragmentation, a prominent neutral loss of phosphoric acid (Δ98 Da) is generally observed, and it has been suggested that a triplet neutral loss fingerprint could occur for specific phosphoresidues, but recently this has been called into question ^5–7^. One of the main reasons why these phosphoramidate bond-containing phosphoamino acids, such as phosphohistidine, have remained unexplored is their instability ^8,9^. The phosphoramidate bonds are acid-labile and thermosensitive, and because most of the phosphoproteomic protocols use acidic conditions (< pH 5) for phosphopeptide enrichment and run LC-MS/MS in acidic buffers (∼pH 2), this means that only the most stable O-phosphate ester phosphoresidues are detected.

Despite rapid advances in MS speed and sensitivity, because of the wide range of stoichiometry of phosphorylation at different sites not every phosphopeptide can be identified without utilizing a method for phosphopeptide enrichment. The development of efficient methods for phosphopeptide enrichment has been essential to the rapid progress in phosphoproteomics ^10^. The most common methods used to enrich phosphoproteins and phosphopeptides without any specificity for sequence motifs or phosphoresidue are iron- or gallium-IMAC (Immobilized Metal Affinity Chromatography) and MOAC (Metal Oxide Affinity Chromatography), such as titanium dioxide, which rely on the affinity of metal ions for phosphate under acidic conditions. Because of the strong binding of phosphate, complete elution of bound phosphopeptides is difficult, and there is also some non-specific binding of acidic peptides. Other chromatography methods, such as HILIC (Hydrophilic Interaction Liquid Chromatography) and ERLIC (Electrostatic Repulsion Hydrophilic Interaction Chromatography) where the effect of pH on analyte charge state varies based on each compound’s pKa, or prefractionation approaches with SCX (Strong Cation Exchange) have been used but are not all optimal for non-acidic pH as they generally need low pH elution. A few methods for phosphate binding at neutral pH exist. Phos-Tag chromatography can bind and elute phosphate at neutral and physiological pH, but is more suited for pure proteins, rather than complex phosphopeptide samples. Hydroxyapatite (HAP) resin has a similar affinity for phosphopeptides under non-acidic conditions but can be applied to complex samples ^11^. The structure of HAP is close to the chemical structure of natural bones or coral and can therefore be used at physiological pH to bind phosphate, although HAP is sensitive to degradation under acid conditions and its use is not advised for standard acid elution ^12^.

Another approach commonly used to enrich phosphopeptides is ImmunoAffinity Purification (IAP) using antibodies, generally crosslinked to a resin, targeting a consensus phosphomotif or a specific phosphoresidue, such as phosphotyrosine (pTyr) ^13,14^. This last option is highly specific but is generally limited due to the cost of use. IAP can be less efficient in complex samples due to the high background that dilutes the target, but this can be offset by combining with pre-fractionation or prior global phosphopeptide enrichment. Following the recent development of phosphohistidine (pHis) monoclonal antibodies (mAbs) ^15^, we have developed a fully non-acidic method using immunoaffinity purification to enrich non-conventional phosphorylation site peptides that is compatible with conventional LC-MS/MS analysis.

## Results

To develop and validate a method for purifying pHis-containing peptides from tryptic digests of a human cancer cell line (HeLa), we used synthetic His-containing peptides chemically phosphorylated with phosphoramidate, confirmed to contain pHis by peptide dot blotting with anti-pHis mAbs, thin-layer chromatography and MS/MS analysis. This purification consists of a strict non-acidic method compatible with hydroxyapatite (HAP) or strong-anion exchange chromatography (SAX) for global enrichment of all phosphopeptides and phosphoresidues, under conditions designed to conserve pHis, pArg and pLys. A graphical schematic of the different experimental approaches we tested for phosphopeptide enrichment under non-acidic conditions is illustrated (Fig. 1). Briefly, an immunoaffinity purification step using 1- and 3-pHis mAbs crosslinked to protein A resin was carried out either prior to or after global phosphopeptide enrichment using HAP or SAX. pHis-enriched tryptic peptides were then analyzed by LC-MS/MS with CID fragmentation on an LTQ Velos Orbitrap, or HCD fragmentation on a Q-Exactive, and identified using a specific configuration of MaxQuant software, simultaneously considering the detection of Ser, Thr, Tyr, His, Lys, and Arg phosphoresidues with a 98 Da neutral loss and exclusion of C-terminal phosphorylation, since phosphorylation of Lys and Arg strongly decreases trypsin cleavage. Moreover, this could potentially redirect the protease to another close residue for alternative cleavage, and explain the increase of missed cleavages in non-acidic conditions ^16^. All the phosphoresidue identifications from MaxQuant analyses, obtained by different methods corresponding to HAP, SAX and/or pHis enrichment (IAP) under non-acidic conditions, were combined using the software Perseus into a single non-redundant list.

**Fig. 1:**
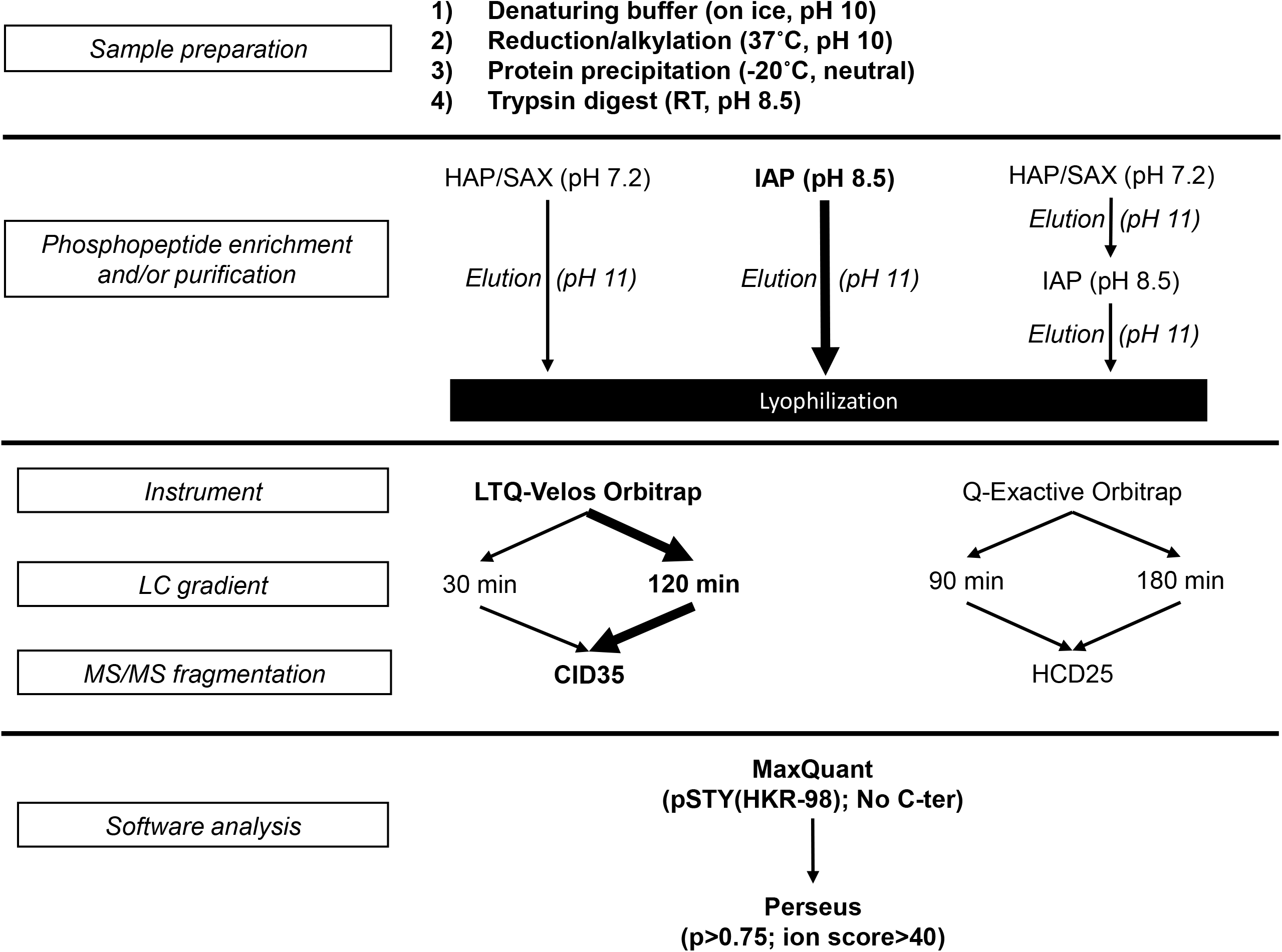
Different options for enrichment and analysis were tested. The most straightforward and main method used is highlighted in bold.

### Synthetic peptides chemically phosphorylated on histidine are enriched and detected by mass spectrometry

Relying on previous work showing that phosphoramidate preferentially phosphorylates histidine ^17,18^, chemically phosphorylated synthetic peptides were chosen as suitable tools for optimization and validation of our method and protocol. Phosphoramidate was synthesized according to Wei and Matthews ^19^, and its ability to chemically phosphorylate synthetic poly-histidine (P9386, from Sigma) was confirmed by two methods: in-gel staining (using Pro-Q Diamond) and blotting with our 1/3-pHis monoclonal antibodies (Supplementary Fig. 1A-B). The N-phosphate chemical phosphorylation of recombinant protein histone H4 was also revealed by dot blotting using our 1/3-pHis mAbs (Supplementary Fig. 1C). In addition, Thin-Layer Electrophoresis (TLE) at pH 8.9 revealed the phosphorylation of a synthetic peptide from histone H4 (Supplementary Fig. 1D) containing the known H18 site (GAKR**<u>H</u>**RKVL), but lacking “conventional” Ser, Thr or Tyr phosphosites.

The NME1 (H118) and PGAM1 (H11) enzymes are phosphorylated on their active site His, forming 1-pHis and 3-pHis isomers, respectively. To test our experimental enrichment conditions a panel of three peptide sequences were synthesized to generate known 1/3-pHis containing peptides: the NME1 H118 site (RNII**<u>H</u>**GSDS-NH2), the PGAM1 H11 site (VLIR**<u>H</u>**GESA-NH2) and the histone H4 H18 site (GAKR**<u>H</u>**RKVL-NH2). Following chemical phosphorylation, each peptide was directly injected into an LTQ-Velos Orbitrap for analysis; a manual annotation of the MS spectrum for histone H4 phosphopeptide is illustrated (Fig. 2).

**Fig. 2:**
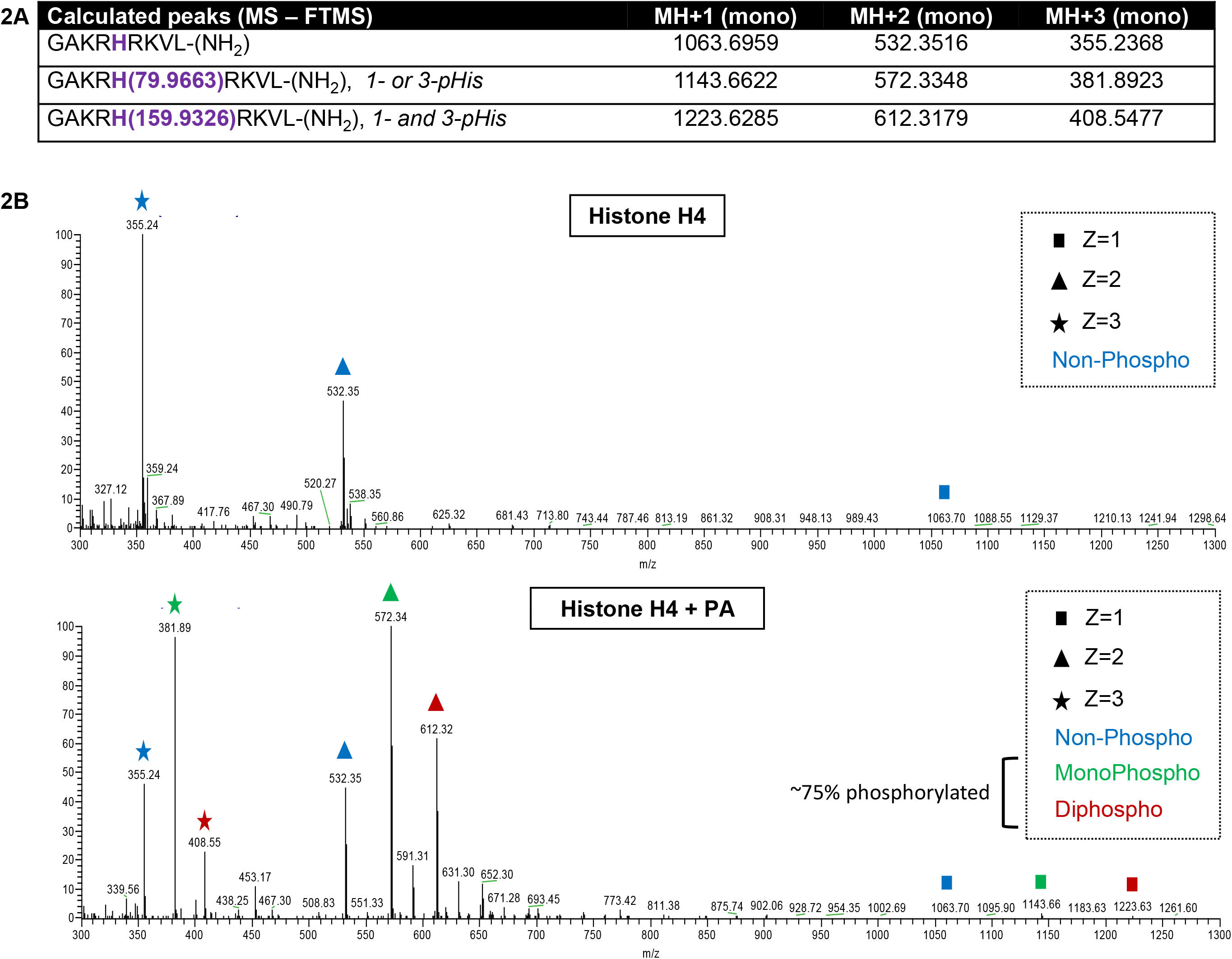
MS detection of pHis after direct-injection of chemically phosphorylated histidine-containing synthetic peptides chemically phosphorylated. **A**. Theoretical masses of histone H4 peptides and its 1/3-pHis containing forms (using MS-product from Protein Prospector V5.20.0 - UCSF). **B**. After single-injection on a LTQ-Velos Orbitrap mass spectrometer (CID fragmentation), manual observation of MS profile using Xcalibur software allows detection of peaks corresponding to non-phosphorylated (blue), monophosphorylated (green) and diphosphosphorylated (red) forms of histone H4 peptides. (Similar observations were made for chemically phosphorylated NME1 and PGAM1 peptides - data not shown). PA: Phosphoramidate. The charge state (z) is symbolized by a square (+1), triangle (+2) or a star (+3).

In addition, a time-course experiment conducted using an autosampler to inject a pool of chemically phosphorylated peptides (NME1, histone H4, PGAM1) every 30 min with a standard pH 2 buffer for MS/MS, revealed that chemical phosphorylation persisted after more than 50 h when the reaction was kept at 4°C in 100 mM TEAB (Triethylammonium bicarbonate) buffer, pH 8. New chemical phosphorylation due to remaining free phosphoramidate should be strongly decreased at 4°C ^19^. *In silico* analysis, using a decoy enzyme on MaxQuant to identify the correct masses of the synthetic peptides with standard database searching, allows automated detection of His phosphorylation (Supplementary Fig. 2). The pHis-containing peptides could be identified for each sequence, suggesting that at pH 8, the phosphoramidates should be stable long enough to allow enrichment. Furthermore, a peak corresponding to a diphosphorylated form of the histone H4 peptide was detected, suggesting that the presence of either a 1/3-diphospho-histone H4 peptide or at least two phosphoramidate bonds, one of which would have to be either pLys or pArg, since the peptide lacks Ser/Thr/Tyr residues. Based on this observation, the potential phosphorylation of N-phosphate residues, Lys and Arg, were also considered in our subsequent search.

To compare the profile of the same sample after IAP enrichment alone or combined with global phosphopeptide enrichment, the three peptide sequences were separately phosphorylated with phosphoramidate and then pooled. After non-acidic pHis enrichment using separate or successive hydroxyapatite (HAP) and 1/3-pHis immunoaffinity purification (IAP), nLC-MS/MS analysis was done using a NanoSpray Ionization (NSI) source on a LTQ-Velos Orbitrap ^20^ to follow the stability and purification of pHis through the process. The different liquid chromatography profiles showed that IAP alone, or successive HAP and IAP, specifically decreased the amount of total peptide material, as indicated by the reduced complexity of the sample (Supplementary Fig. 3A). Subsequent manual analysis of the MS/MS spectra of the dual-enrichment sample (Supplementary Fig. 3B-C) showed that phosphorylated forms of each peptide were still detectable in their three different charge states (z).

### Strict non-acidic conditions permit purification of pHis containing peptides from human cancer cell lines

To assess the stability of the natural phosphorylation profile of His, Ser/Thr and Tyr residues (Fig. 3A), during the steps used to prepare the phosphopeptides from human cells prior to enrichment, we carried out immunoblotting experiments as follows: a mix of 1/3-pHis mAbs (SC1-1 and SC39-6), the 4G10 pTyr mAb, and a pAkt-substrate (RxxS/T) mAb were independently used to monitor, His, Tyr and Ser/Thr phosphorylation, respectively. The immunoblot revealed that denaturing lysis of cells in 8 M urea buffer (pH 10), reduction with DTT (pH 10) and alkylation with chloroacetamide (pH 10), followed by protein precipitation with cold methanol, all carried out under non-acidic conditions were compatible with the global stability of histidine phosphorylation on proteins from human cells (HeLa and ALVA-31) (Fig. 3B). After overnight trypsin digestion at pH 8.5, a peptide dot-blot revealed that 1-pHis and 3-pHis signals were still conserved on phosphopeptides (Fig. 3C).

**Fig. 3:**
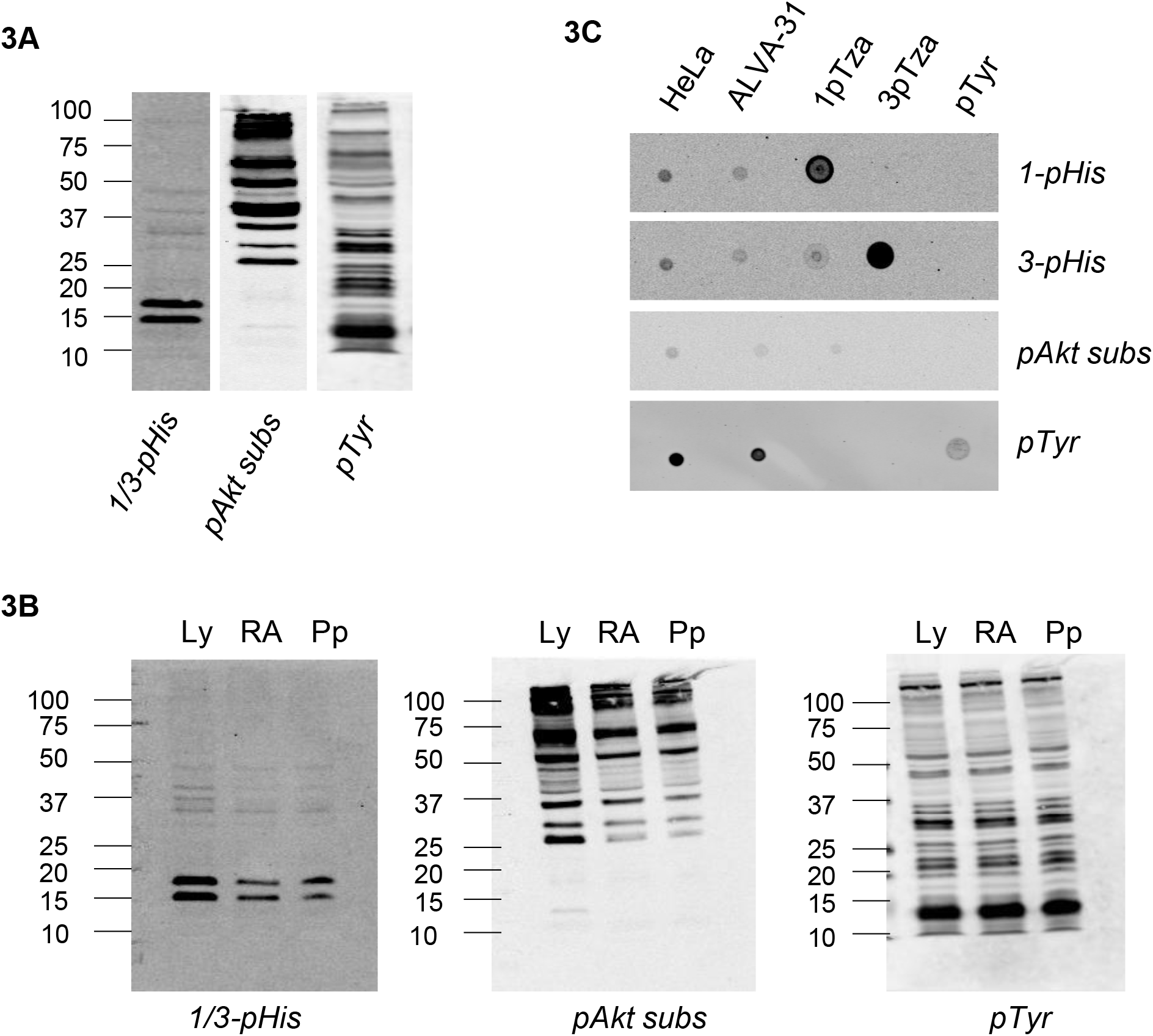
Phosphopeptides preparation under non-acidic conditions from human HeLa and ALVA-31 cell lines. **A-** Phosphorylation profile of HeLa lysate in 8 M Urea at pH 10 run in triplicate on the same gel to an adjust pHis, pSer/Thr and pTyr in parallel using, respectively, a mix of 1/3-pHis mAbs (SC1-1, SC56-2), pAkt substrate (RxxS*/T*) mAb (pAkt subs) and 4G10 pTyr mAb. **B-** Phosphoprotein extraction in non-acidic/neutral condition. Ly: Lysate (Urea 8 M, pH 10), RA: Reduction/Alkylation step, Pp: Protein precipitation (MetOH). **C-** Peptide dot blot after trypsin digest, pH8.5. The 1pTza (phosphoryl-triazolylalanine) and 3pTza peptide libraries correspond to randomized sequences of non-hydrolyzable 1- and 3-phosphohistidine analogues.

Several different approaches were tested to develop the best method for enriching and preserving phosphoramidate bonds in tryptic digests of proteins from confluent HeLa cell lysates (Fig. 1). Global phosphopeptide MS analyses were made with an LTQ-Velos Orbitrap using Collision Induced Dissociation (CID) or with a Q-Exactive Orbitrap using High Collision Dissociation (HCD). In figure 4, the proportion of each phosphoresidue is indicated for each enrichment or purification method, independently or in combination. Only the highest probability phosphate localizations (p>0.75) were considered. Global phosphopeptide enrichment with HAP or SAX (Fig. 4A-B) under non-acidic conditions revealed that a median coverage proportion of ∼52.5% for phosphoramidate (pHis, pLys and pArg) was obtained for the two methods, which, surprisingly, was greater than for conventional pSer, pThr, pTyr phosphoester bonds (∼47.5%). The median of pHis peptides was 5.5%, which corresponds to the expected ratio if we consider a deterministic model of the complete human proteome where all 9 potential SONAte phosphoresidues would be phosphorylated (Supplementary Fig. 4A-B). For IAP, we used of a mixture of three specific anti-1-pHis mAbs (SC1-1, SC50-3, SC77-11) and three specific 3-pHis mAbs (SC39-6, SC44-1, SC56-2) coupled to protein A resin (Fig. 4C). Through using six mAbs we hoped to capture both 1-pHis and 3-pHis residues embedded in as wide a variety of primary sequence contexts as possible. Using IAP alone, the proportion of phosphoramidate-containing peptides after enrichment increased up to 80% of the recovered phosphopeptides, with 23% significant pHis site localizations. Interestingly, a significant proportion of pArg and pLys were detected with this method. However, the successive combination of both global phosphopeptide enrichment (using HAP or SAX) and pHis purification that should help to increase specificity, resulted in lower overall efficiency than IAP alone (Fig. 4D-E). This suggests that even if the HAP/SAX plus IAP combination is better than HAP or SAX alone, the additional steps and longer experimentation time most probably led to a decrease in recovery of non-conventional phosphopeptides during enrichment due to their chemical lability. When multiple phosphorylations occur within the same peptides, peptides with conventional and non-conventional phosphates can co-elute, meaning that the enrichment cannot be totally specific and justifying why other phosphoresidue options should be considered. For this reason, the proportion of monophosphorylated and multiphosphorylated peptides, for phosphoresidues with a high probability of localization, is also indicated for each type of enrichment (Supplementary Fig. 5); only the identity of the mapped phosphosite is considered.

**Fig. 4:**
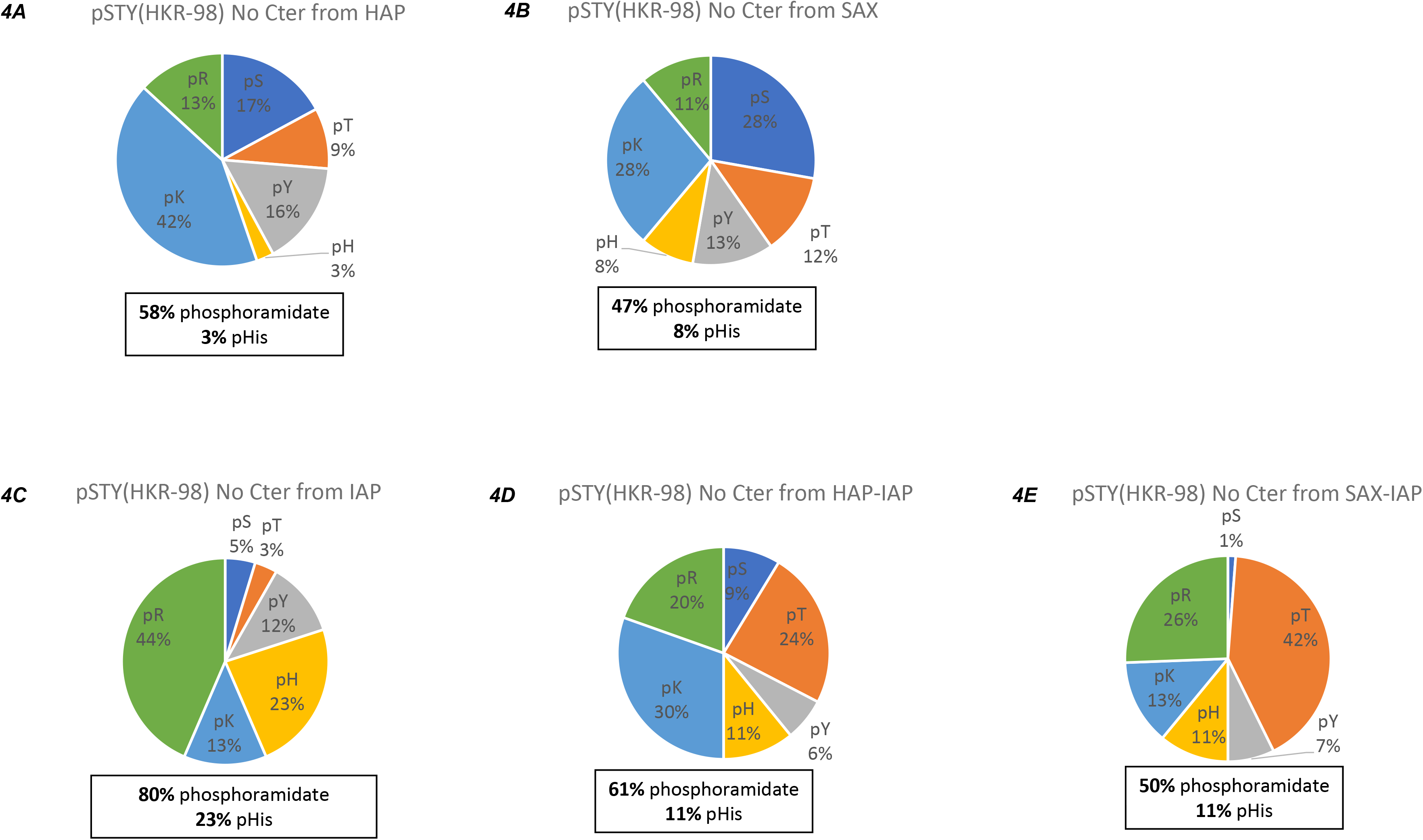
Proportion of phosphoresidues after phosphoenrichment. Values indicated correspond to the number of phosphorylation sites identified by MS/MS and their percentage per residue. Only O-phosphate (STY) and N-phosphate (HKR) residues were integrated in MaxQuant analysis for each enrichment method, considering the main neutral loss of 98 Da for the N-phosphate residue. Only phosphorylation sites identified with a probability of localization >0.75 are shown. No phosphorylation in C-ter position were considered. **A. and B**. Global phosphopeptide enrichment from HAP (hydroxyapatite) or SAX (strong anion-exchange) chromatography using the same lysate sample. **C-** Unique IAP enrichment using 6 combined pHis mAbs (1-pHis clones: SC1-1, SC50-3, SC77-11; 3-pHis clones: SC39-6, SC44-1, SC56-2). **D. and E**. Combination of global phosphopeptide enrichment, followed by 1/3-pHis immunoaffinity purification using the same lysate sample.

Combining multiple samples and MS/MS runs for every method, a list of 32,055 unique peptides for 5,092 unique proteins was extracted from raw data (Fig. 5A). Combining all phosphopeptide enrichment methods, 8,694 histidine containing peptides were purified with 5,347 identified in an IAP experiment using pHis mAbs. 36 of these histidine containing peptides were previously reported to be phosphorylated on a specific His. Indeed, only a few pHis sites are known on mammalian proteins (∼30), and some of these are in tryptic pHis-containing peptides that are too short or too long to be considered by mass spectrometry using our parameters, like the histone H4 (H18) and ACLY (H760) peptides, respectively. After fragmentation with a cut-off value at 40 for ion score, reflecting a high confidence measure of how well the observed MS/MS spectrum matches the predicted spectrum of the stated peptide, over 2,567 phosphorylation sites were considered. 1,476 non-redundant phosphorylation sites were selected based on a probability of localization higher than 75% with a median of 95.3%. Among them, 425 phosphosites correspond to new phosphoramidate bonds (Supplementary Table 1) with GO molecular functions related to binding to nucleoside phosphates, ions and heterocyclic compounds, as well as being involved in protein dimerization activity.

**Fig. 5:**
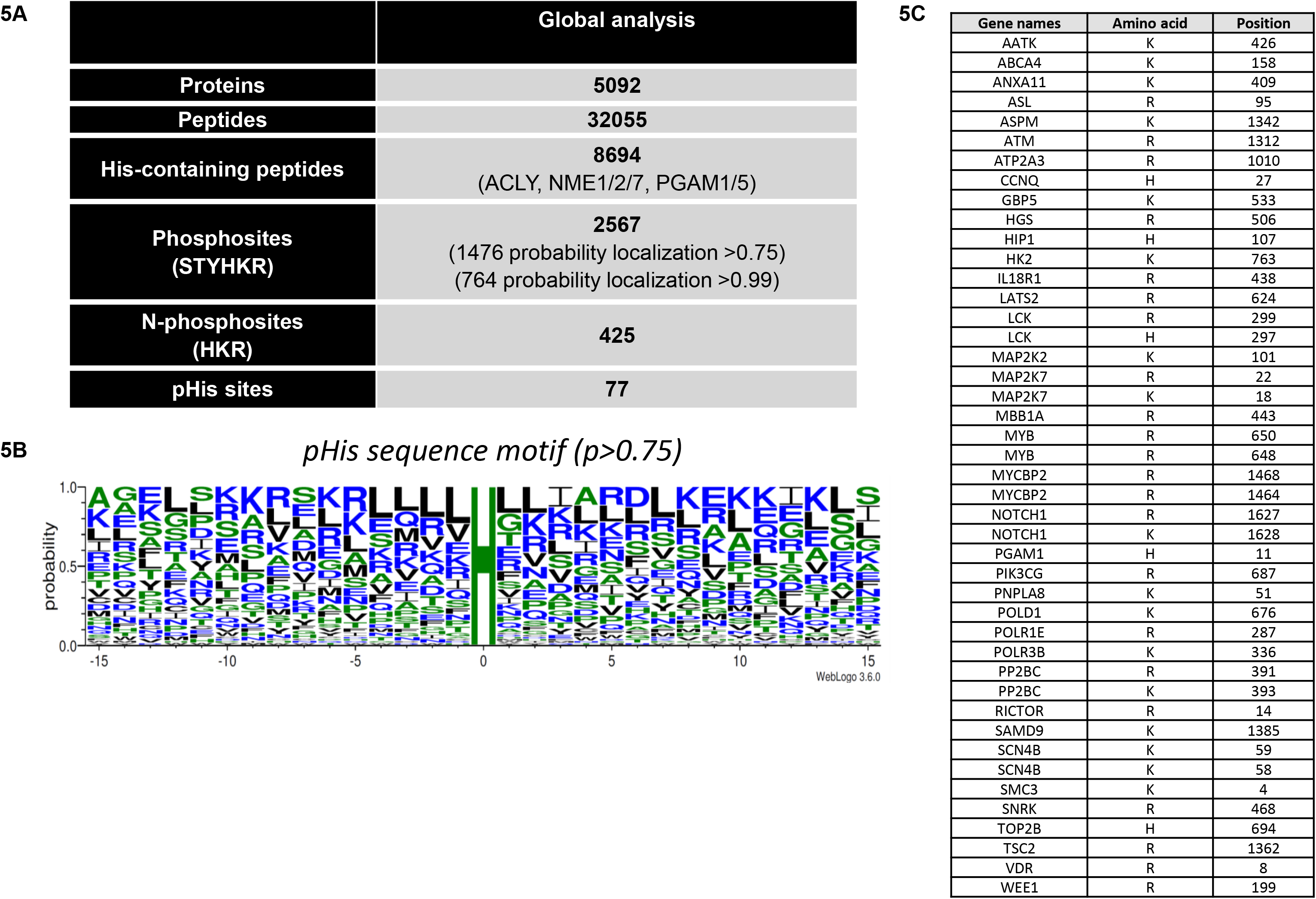
Phosphohistidine sites identification from human HeLa cells. **A**. Combined global identification from all non-acidic enrichment methods, using MaxQuant software and Perseus, confirmed the known phosphorylation 3pHis site (H11) of PGAM1, with a strong residue probability of 99.95% (data not shown). **B**. Sequence motif extracted from the significant phosphohistidine containing peptides list (p>0.75) using the sequence generator WebLogo. Residues in blue, green and black represent, respectively, the hydrophilic, neutral and hydrophobic residue categories. **C**. Selection of N-phosphate phosphoresidues (from the complete list in Sup. Table 1) and their position, with probabilities of localization higher than 95% and without identification of close pSer, pThr or pTyr possibilities.

Seventy-seven of these phosphorylation sites were localized to histidine residues, including the known active site H11 of PGAM1. Physiological phosphorylation of this site was confirmed on the MS/MS spectrum by the b/y ion series (Supplementary Fig. 6) extracted on the Integrated Proteomic Pipeline IP2 ^21^. In addition, in a separate search considering pHis immonium ions, pHis 11 of PGAM1 was also identified with a diagnostic peak (data not shown). No major sequence motif was defined from the sequence window of these 77 pHis peptides using Weblogo 3 ^22^, but Leu and Ile were overrepresented around the pHis residue (Fig. 5B). Interestingly, Leu residues were globally overrepresented in the sequence motifs surrounding the three phosphoramidate bonds on His, Lys, and Arg but this was not the case when considering the phosphopeptides with phosphorylated Ser, Thr, and Tyr (data not shown). Similar observations were also made by Hardman et al. ^5^ and might suggest selectivity for phosphoramidates bonds or a preferential adsorption to the resins used, but most probably stabilization against hydrolysis by local hydrophobic acids.

## Discussion

Conventional phosphorylation is, by definition, confined to the three O-phosphate bonds (pSer, pThr, pTyr). Non-conventional phosphorylation includes the three N-phosphates (pHis, pLys, pArg), two acyl-phosphates (pAsp, pGlu) and S-phosphate (pCys). Several interesting proteins with non-conventional N-phosphorylation sites were identified through the non-acidic enrichment protocol we have tested (Fig. 5C). Further analysis of these sites will be required for validation of their identity and to determine whether they are of functional significance. Some of these sites occur in well-studied protein kinases, like the Lck tyrosine kinase (H297 and R299 – both in the catalytic domain) and the ATM DNA damage response kinase (R1312), and these phosphorylations could regulate their signaling output. Other novel N-phosphorylation sites include TOP2B (H694) and POLD1 (K676) involved in DNA transcription and replication; TSC2 (R14) and Rictor (R1362) in the mTORC1 and mTORC2 pathways, respectively, that regulate cell growth and respond to nutrient status and growth factor signaling; and MYCBP2 (R1464 and R1468), recently identified as the single member atypical Thr-specific E3 ubiquitin-protein ligase ^23^.

At this stage, the stoichiometry of His, Lys and Arg phosphorylation is not known but considering the increase of phosphoramidate bonds using IAP with six pHis mAbs, it seems likely that we are most successful in enriching the most abundant pHis, pLys and pArg tryptic peptides, even if there may be also some primary sequence bias in recovery. Somewhat surprisingly, a significant number of these pLys and pArg-containing peptides were stable and identified through our IAP non-acidic global phosphopeptide enrichment protocol. Within the human proteome, Lys and Arg are in total ∼4 times more common than His (Supplementary Fig. 4A), which would increase the chances of detecting these phosphorylated residues, if phosphorylation frequency is proportional to abundance, and all three N-phosphates have the same phosphoramidate stability. Although our original characterization of the anti-1/3-pHis mAbs demonstrated that they did not recognize pSer, pThr and pTyr residues, the observed enrichment of pLys and pArg peptides suggests that under IAP enrichment conditions, where a mixture of pHis mAbs is used at very high density on the beads, these mAbs may also recognize phosphoramidate linkages per se. Further studies are underway to obtain a structural understanding. In our prior SILAC-based studies, 687 potential pHis-containing proteins were identified by using IAP, with the 1-pHis mAbs SC1-1 and the 3-pHis mAbs SC39-4, on proteins from denatured 293T cell lysates ^15^. His-containing peptides from 579 of these proteins were also identified among the non-acidically-enriched phosphopeptides, with 17 phosphoramidate sites enriched and identified by MS. Among these phosphoramidate sites, 5 were identified as pHis: FBL (H93, possibly 3-pHis), PGAM1 (H11), SSRP1 (H449, possibly 1-pHis), TOP2B (H694), ZNF638 (H527, possibly 1-pHis). PGAM1 is known to contain exclusively 3-pHis and was isolated from a 3-pHis IAP experiment, but since we used a combination of 1- and 3-pHis mAbs to increase coverage of enrichment, we currently cannot determine if any of the other identified pHis sites are isomer specific.

Whatever the method used, our data indicate that global phosphopeptide enrichment under non-acidic conditions allows enrichment of phosphopeptides with non-conventional linkages. Acidification of a samples strongly decreased the proportion of non-conventional phosphoresidues like pHis recovered from HAP chromatography (Supplementary Fig. 7). However, even using non-acidic conditions, we noted that several His present in the 8,694 His-containing peptides that were enriched or purified have been reported to be phosphorylated, and more than likely they were enriched as a result of their phosphorylation (CCT7 H346, GAPDH H111, HIST1H4A H75, LDHA/B H67, NME1/2 H118, PGAM1 H11, RPS3A H232, SUCLG1 H299, TUBB H105), but their phosphate was then lost during LC-MS/MS analysis due to the acidic conditions, with in the end only 77 pHis sites being detected and localized with high probability (greater than 0.75 with a median of 0.953). Of course, the presence of His in an enriched peptide is not enough to prove that it was phosphorylated, but there seems little doubt that the positive ionization parameter, which requires highly acid (pH 2-3) chromatography, remains a limiting step in the identification of additional non-conventional phosphorylation sites that were in fact enriched. As mentioned by Sickman and Meyer, the half-life life of pHis is about 30 min at pH 3 ^9^. But a higher pH, like pH 5, tends to strongly reduce ionization efficiency and thereby increase peptide loss at the MS stage (Supplementary Fig. 8). Alongside the development of our method using pHis mAbs, other recent efforts have been made to enrich and reveal His phosphorylation events by MS. For instance, Potel et al. generated a phosphoproteome from *E*. *coli* recovering 10% pHis sites (considering only pS/T/Y/H possibilities) using Fe^3+^-IMAC and MS/MS analysis with HCD fragmentation before MaxQuant identification ^24^. Progress employing high-pH separations and negative electrospray ionization to generate neutral loss were also made, as well as development of new fragmentation methods like the UVPD (Ultra-violet PhotoDissociation) ^25,26^. However, additional efforts by the proteomics research community are required to define the best resolution and fragmentation methods for non-conventional phosphosites and develop new technology compatible with identification of the full range of phosphorylation events. In this regard, the electron transfer/higher energy collisional dissociation (EThcD) has recently been reported to fragment doubly-charged phosphopeptides without loss of the labile phosphates on Arg, His, Cys, and Lys as well as Ser residues using synthetic peptides ^27^. In addition, the development of nETD (negative ion mode) fragmentation and MS analysis, where the polarity is reversed, should be highly compatible with a non-acidic phosphoproteomic analysis ^28^.

Interestingly, the chemically phosphorylated synthetic peptides we analyzed by direct injection illustrate that short exposure to acid buffer during fast chromatography is recommended, as only 25% of the peptides are not phosphorylated, which means that at least 75% of the chemical phosphoramidate bonds remained stable through the positive ionization (Fig. 2B). Considering that these 25% non-phosphorylated peptides could possibly come from a partially incomplete chemical phosphorylation. But even after pHis enrichment and then removal of non-chemically phosphorylated peptides, a 30 min chromatography induced about 50% degradation of these acid-labile phosphorylations (Supplementary Fig. 3B). Nevertheless, the time for liquid chromatography is also dependent on the complexity of the samples and must be adapted and considered for the accuracy and reproducibility of the identification, if sample comparison is required. An immediate solution would be to increase the length of the C18 column while increasing the flow rate, to the detriment of pressure increase within the column, in order to reduce a time-dependent exposure to the acid gradient. Another option remains the use of non-acidic fractionation. Fast protein fractionation from the PC-3 human prostate cancer cell line under non-acidic conditions (90% MetOH) revealed the possibility of conserving, detecting and fractionation pHis-containing proteins (Supplementary Fig. 9A). Considering that most of these proteins are phosphorylated on their histidine, family members of the known pHis-containing NME and PGAM proteins were mainly collected in the late fraction and identified by mass spectrometry (Supplementary Fig. 9B).

Besides improved stability and fractionation to make pHis peptides more abundant during MS/MS analysis, a way to increase the non-conventional phosphorylation site identification is to reduce the low localization probability due to proximity of any potential phosphoresidues. As shown with the chemically phosphorylated histone H4 peptide (Supplementary Fig. 2), the detection of a phosphoramidate bond is undeniable, but the proximity of other N-phosphates residues (Arg and Lys), makes it statistically more difficult to get a perfect ion series coverage. A similar observation was made in our enrichment analysis with the identification of phosphoramidate bonds but technical difficulties in precise assignment of the exact position (Supplementary Table 2). Efficient compatible fragmentation or additional NMR validation would be an asset to better define the exact localization.

Based on earlier studies it has been suggested that pHis might be more abundant in proteins than expected ^29^ and the existence of His kinases and pHis phosphatases in human cells also implies the existence of non-conventional phosphorylation-based signaling systems ^30–33^. Our work shows for the first time that under non-acidic conditions, an overall coverage of SONAte phosphopeptide enrichment using HAP can reveal more non-conventional than conventional phosphorylation sites (Fig. 4A and Supplementary Fig. 4B) and that phosphopeptides with phosphoramidate linkages can be effectively enriched using immobilized pHis mAbs for IAP (Fig. 4C). A truly global phosphopeptide enrichment would require the inclusion of non-conventional pAsp, pGlu and pCys phosphosites. The complete SONAte experimental consideration made in the supplementary data 4B from HAP enrichment suggests that pAsp and pGlu represent respectively 11% and 16% of the phosphoproteome under non-acidic conditions whereas pCys was only 2%. These observations give a new impetus for the study of the still hidden phosphoproteome in multicellular organisms.

Our reported number of pSer/pThr/pTyr sites with a probability of localization higher than 75% (1051) was significantly lower than expected from published databases of HeLa cell phosphosites enrichment (5000-8000). We do not fully understand the reason for this, but it suggests that the non-acidic conditions are not optimal for phosphate-ester enrichment. First, pSer/pThr levels can be moderately decreased during alkaline treatment of proteins and peptides (Fig. 3B-3C using pAkt substrate). Moreover, pAkt-substrate antibody motif recognition (RxxS*/T*) may be reduced after digestion, because of cleavage after the Arg. Second, the simultaneous consideration of additional phosphoresidues increases the number of search possibilities, which decreases the probability of localization “by default” on Ser/Thr/Tyr residues, and paradoxically can also decrease below 0.75 the probability of true localization, that would then be excluded by our filter if the ion series are not able to distinguish clearly the position of the phosphate between two close residues. Third, the condition of enrichment or elution itself could reduce the binding efficiency of O-phosphate-containing peptides, which could explain why many peptides with more stable multiple phosphorylations were enriched, as a result of a higher avidity for HAP/SAX. But interestingly, NME1 pSer120, NME7 pThr304 and PGAM1 pSer14 peptides were also enriched and identified after the successive HAP-IAP.

Histidine kinases and phosphatases are physiologically important. In bacteria, with function being mainly in the well-known two-component sensor relays. The human pHis proteins we identified include nuclear and cytoplasmic proteins, but few membrane pHis proteins (e.g. ion channels), where recovery is usually poor. The identities of the pHis proteins we identified suggest that pHis may play a role in mammalian cell signaling ^32–34^. Indeed, a direct correlation of pHis protein levels with carcinogenesis has recently been reported ^30,35–37^. Interestingly, we identified 20 N-phosphate sites on proteins that were reported to be enriched by IAP by Hindupur et al. from mTOR-driven hepatocellular carcinoma mouse tumors as potential LHPP substrates, including BCLAF1, which in our studies was found to contain pLys (K391) with a reasonable probability of localization of 79.6%. The use of different experimental approaches and enrichment methods like the one we introduced here, in combination with more traditional conditions ^24,38^, or the development of additional cost-effective specific phosphopeptide binders, like the Src homology 2 (SH2)-domain-derived pTyr superbinder ^39^, would, together with nETD MS technology, help to accelerate progress in the largely unexplored field of non-conventional phosphorylation of proteins.

## Methods

### Synthesis of peptides

The following peptides: NH2-Arg-Asn-Ile-Ile-His-Gly-Ser-Asp-Ser-NH_2_ (NME1 H118); NH2-Val-Leu-Ile-Arg-His-Gly-Glu-Ser-Ala-NH_2_ (PGAM H11); NH2-Gly-Ala-Lys-Arg-His-Arg-Lys-Val-Leu-NH_2_ (histone H4 H18); were synthesized manually, as previously described ^15^, by Nα-Fmoc methodology on PS-Rink resin. The final peptide was >95% pure as determined by tandem analytical LC-MS. The structure of the pure peptide was confirmed by high-resolution electrospray ionization mass spectrometry (ES-MS). Other peptides from library were synthesized in-house on a Protein Technologies (Gyros) “Tribute” peptide synthesizer equipped with real time UV monitoring, using standard Fmoc chemistry.

### Chemical phosphorylation

Phosphoramidate was made according to the method of Wei and Matthews. Peptides were resuspended in water at 10 mg/mL and phosphoramidate resuspended in water at 81 mg/mL. For each reaction, one volume of peptide was added separately to four volumes of phosphoramidate solution. The final concentrations were 2 mg/mL and 65 mg/mL, corresponding to a ratio of 1 mole of peptide reacted with ∼350 moles of phosphoramidate. The reaction proceeded in water for 2 h at room temperature. The completed reaction can be stored at 4°C or immediately diluted 10X with Buffer A (5% ACN, 95% H2O, 0.1% formic acid, pH 2) for direct injection onto the mass spectrometer.

### Immunoblotting

For immunoblotting: Buffers were adjusted to pH 8-9 to stabilize pHis, and methods were modified to avoid heating the samples. The protein samples were prepared in pH 8.8 sample buffer (2% SDS, 50 mM Tris-HCl, pH 8.8, 0.004% bromophenol blue, 10% glycerol, 10 mM EDTA, 100 mM DTT) for electrophoresis. Cell lysates were prepared by rinsing 70-100% confluent 10 cm^2^ dishes twice with 5 ml cold PBS buffer (PBS -/-, pH 8). Cells were scraped directly into 0.5 mL of 2X pH 8.8 sample buffer, incubated on ice and a tip sonicator was used (3x 10 sec bursts/10 sec ice). A syringe and needle (18-20G) can also be used to fragment DNA and tissue as a first step, but sonication is still recommended to limit persistent protein complexes as boiling is prohibited. Lysates were clarified by centrifugation (14,000 x g for 10 min at 4°C) and analyzed immediately using freshly prepared Bis-Tris polyacrylamide minigels with a modified, pH 8.8 stacking gel and either 10% or 12.5% resolving gels. For better stability, the polymerized acrylamide gel can be placed at 4°C 15-20 min before to load samples. The Running Buffer (1X 20 L, pH 8.5: 20 g SDS, 60 g Trizma Base, 288 g glycine, dH2O to 20L) as well as the Transfer Buffer (1X 4 L, pH 8.5: 56.7 g glycine, 4 g SDS, 12 g Trizma Base, 800 ml MetOH, dH2O to 4 L) were cooled at 4°C overnight before use. All electrophoresis steps were performed in a cold room at 4°C and samples were resolved at 100 V for about 2 h. Proteins were transferred to Immobilon-FL PVDF membranes, previously activated in MetOH for 30 sec and equilibrated in transfer buffer for 5 min, at 30 V for 12-18 h at 4°C or at 75 V for 2 h on ice. If Red Ponceau staining is required, a non-acidic solution in PBS 0.2X is used. The blots are immediately incubated for a minimum 1 h at 4°C in Casein Blocking Buffer (0.1% casein, 0.2X PBS -/-, adjusted to pH 9 with 10 N NaOH). From that solution the pH will be conserved for primary antibodies and secondary antibodies solution. All primary antibodies (anti-pAkt substrate mAb (9611S) from Cell Signaling, pTyr mAb clone 4G10 (Cat. 05-321) from EMD Millipore), were diluted at 0.5 µg/mL in pH 9 blocking buffer with 0.1% Tween-20, incubated with membranes for 1 h at 4°C. Membranes were washed three times for 5 min each with 0.1% TBS-T (adjusted to pH9 with NaOH 10N) at 4°C before incubation with secondary antibodies, diluted 1/20 000 in Blocking buffer pH 9 with 0,1% Tween-20 and 0,01% SDS, for 1h at 4°C. After incubation with secondary antibodies, membranes were washed 3 times for 5 min each with 0.1% TBS-T pH 9 at 4°C. Immunoblots were imaged on a LI-COR Odyssey Infrared Imaging System using both channels of the Odyssey (700 and 800) if necessary.

For peptide dot-blotting: the stability of pHis in tryptic digests of proteins was assessed by peptide dot blotting. Peptides containing non-hydrolyzable analogues of 1-pHis and 3-pHis, 1-pTza (50 ng) and 3-pTza (50 ng), were used as positive controls. Peptides were spotted at 1 µL (∼2 ug synthetic peptides for chemical reaction and ∼10 ug for peptides from cell lysate digest) per dot on nitrocellulose membrane, left to adsorb for 5 min and then totally dried with airflow. The membranes were then incubated with pH 9 blocking buffer before starting the immunostaining steps using primary and secondary antibodies as for pHis immunoblotting method above.

### Gel staining

Pro-Q Diamond staining: Bis-Tris Acrylamide gels containing phosphoproteins were fixed in cold methanol solution for 30 min at 4°C with gentle agitation, and then washed with deionized water for 10 min. The wash is repeated three times, and the phosphoproteins stained by covering the gel with Pro-Q Diamond solution (P33301, from Invitrogen) for at least 1 h. The gel is washed twice with deionized water for 5 min. Staining was revealed using a Typhoon FLA 9500 imaging system with 532 nm excitation wavelength laser (PMT650 – 50 µm resolution).

Coomassie blue staining: Bis-Tris Acrylamide gels containing protein/peptides are washed once with deionized water for 5 min then fixed and stained in Blue Coomassie solution with isopropanol for 1 h with gentle agitation. The stained gel is washed for 20 min, three times, with deionized water. The gel can be scanned immediately or kept overnight in solution. Coomassie blue protein-stained were imaged on an Odyssey^®^ Imager in the 700 nm channel.

### Thin-layer electrophoresis

Peptides were spotted at 40 µg per origin and dried on thin-layer cellulose plates from Merck. Electrophoresis was run in 1% ammonium carbonate pH 8.9 at constant 1 kV for 25 min ^40,41^. Peptides were then visualized by ninhydrin staining.

### Tissue culture

HeLa cells were cultured in DMEM media (4.5 g/L glucose, L-glutamine and sodium pyruvate) complemented with 10% FBS and antibiotics (penicillin-streptomycin). ALVA-31 and PC-3 prostate cancer cells were cultured in RPMI-1640 media with 10% FBS and antibiotics (penicillin-streptomycin). All cells were grown in a 37°C, 5% CO_2_ incubator.

### Rabbit hybridoma culture

The pHis hybridoma cell lines were maintained by culturing in a 37°C, 5% CO2 incubator. As previously reported by Fuhs et al. in 2015, the Growth Medium corresponds to 1X HAT 240E medium; 500 ml RPMI 1640, 40 ml Rabbit Hybridoma Supplement A (Epitomics) containing antibiotic/antimycotic/gentamycin, 55 μM 2-mercaptoethanol and 10% FBS. Briefly, cultures were seeded at 1 × 10^5^ cells/ml and split at 70-80% confluency by aspirating media and replacing with fresh medium. Cell lines were stored in liquid N2 in freezing media (90% FBS, 10% DMSO).

### pHis mAb production and purification

To produce the pHis mAbs (1-pHis mAbs: SC1-1, SC50-3, SC77-11; 3-pHis mAbs: SC39-6, SC44-1, SC56-2), pHis hybridomas were expanded from 10 cm^2^ dishes to T175 flask using Rabbit Hybridoma Growth Medium. Once confluent, cells were collected by centrifugation at 1,200 rpm (∼300 g) for 5 min. Cells were grown in Serum Free Medium (IS-MAB CD chemically defined medium from Irvine Scientific, 1% antibiotic/antimycotic supplement and 1% Glutamax) into 10-layers Corning HyperFlasks for 7-9 days until cell viability was approximately 50%. To harvest antibodies, cells were collected by centrifugation. Cell supernatants were spun again in fresh tubes at 3,000 rpm for an additional 15 min.

To purify the pHis mAbs, protein-A-agarose beads were washed with PBS before overnight incubation at 4°C with 1 g protein-A-agarose beads per 100 ml hybridoma cell supernatant. The protein-A-agarose beads were pelleted at 1,000 rpm for 5 min at 4°C and washed three times with 3 volumes of beads in cold PBS (pH 7.4). Anti-pHis IgGs were eluted with 10 sequential additions of 1 ml Elution Buffer (100 mM glycine, pH 2.8) at RT. Each sequential elution was immediately neutralized with 1.0 M Tris-HCl (pH 8.3). Anti-pHis mAb concentrations were measured by IgG A280 on a Nanodrop and stored at 4°C. Purified mAbs (0.5 < 260:280 ratio < 1) were used at a concentration of 0.5 μg/ml and validated by immunoblotting cell lysates and dot blotting synthetic peptides.

### Antibody crosslinking to protein A agarose with BS3

Pure antibodies from elution (IgG 260:280 ratio < 1) were dialyzed overnight at 4°C in 1 L PBS -/- 1X using 3,500-10,000 Molecular Weight Cut-Off (MWCO) cassette to remove primary amines, like Tris or glycine, as the crosslinker NHS-ester react with primary amino-groups (-NH_2_). Before crosslinking the pHis mAbs at 1 mg/mL, the protein A beads are washed with 20 mM Hepes, and pooled mAbs added to protein A beads (w/w) in 20 mM Hepes buffer. BS^3^ (bis[sulfosuccinimidyl] suberate, (21580) from Thermo Scientific) is added to a final concentration of 2.5 mM and incubated for 45 min at RT on a rotating wheel. The reaction is quenched by adding 50 mM Tris-HCl pH 7.5 for 15 min at RT on the wheel. Next, the beads are placed into a column and washed three times with 1 bead volume with PBS before storing at 4°C until use. A rabbit IgG column can be prepared in the same way for preclearing or as negative control.

### Cell lysis and peptides preparation

Cells were washed twice on ice with cold PBS-/-(stored at 4°C) to eliminate BSA. Just before use, the phosphatase inhibitors (PhosStop tabs from Roche), Octyl-B-D-Glucopyranoside 30 mM and NH4HCO3 1M pH 8.5 were added to the denaturing buffer (urea 8 M, Hepes 20 mM, pH 10 using NaOH 10N). Make fresh denaturing buffer to limit degradation of urea which can induce carbamylation of proteins. Cells were solubilized at ∼50.10^6^ cells/mL with room temperature denaturing buffer scrapping the cells on ice. Collected cells were vortexed and disrupted on ice passing the lysate through 20G syringe (3-5 times), then sonicated in an adapted tube to shear DNA (3-5 periods of 10 sec burst/10 sec ice). We recommend the use of a Tip sonicator with low intensity ∼2 Watts. In function of the volume, it should not exceed 40 Joules on ice per period or decrease the sonication time to limit additional phosphoramidate degradation. Before any centrifugation, 500 U/mL of benzonase was added to the lysate and supplemented with 1 mM MgCl_2_ and 10 mM DTT for reduction at 37°C for 1 h, then cooled down to RT. Then, 10 mM of freshly made chloroacetamide (C0267, from Sigma) was added to the lysate and incubated at RT for 15 min in dark. Finally, the lysate was centrifuged at 4000 g for 15 min at RT. During these steps, the alkaline condition (pH 10), stabilized the phosphoramidate bonds at moderate temperatures (10°C-37°C). This temperature range is high enough to avoid any urea crystallization because at this concentration (8 M ∼50% w/v) it starts to precipitate below 10°C, and because at this pH urea degradation is more temperature sensitive.

Proteins from the supernatants were precipitated with at least 8 volumes of cold MetOH to one volume of sample supplemented with one volume of Chloroform. The solution was fully vortexed and incubated for a minimum 1 h at −20°C, before spinning down the precipitate at >7500 g for 30 min at 4°C.

Impurities were removed, and the tube and pellet were washed with suspension buffer: 20 mM Tris-HCl pH 8.5 if HAP/SAX enrichment was used first, or immediately with 50 mM NH_4_HCO_3_ pH 8.5 if IAP alone is being used. The wash was removed, and pellet was resuspended in suspension buffer, with the phosphatase inhibitor mix (PhoStop from Roche) added just before use. At this step, the same volume used for cell solubilization (∼50.10^6^ cells/mL) was used to resuspend with a cut pipette tip before sonication if necessary. The protein concentration was immediately measured with a Bio-Rad DC protein assay, adjusting the volume to ensure that the concentration was less than ∼10 mg/mL protein. A few μL were kept aside for control on acrylamide gel or protein dot blot. For pHis staining using the mAbs, use the sample buffer pH 8.8 as recommended for western-blotting. Proteins were digested for at least 14 h (overnight) with an animal-free recombinant bovine trypsin expressed in corn (Trypzean T3568 from Sigma) at 1/50 w/w and RT. Samples were centrifuged 5 min at 20,000 g and supernatant kept at 4°C. The protein digest was checked on 10% acrylamide minigel by blue Coomassie staining. In the meantime, the pHis stability on peptides can be checked directly by dot blot.

Negative control sample for comparative analysis of HAP enrichment were acidified at pH 3 using 0.01% formic acid and boiled at 90°C for 15 min. The pH was neutralized adding 20 mM Tris-HCl pH 8.5.

### Global phosphopeptides enrichment (HAP or SAX)

Chromatography column were prepared using HAP resin (Bio-Rad, 1300420) or SAX resin (Poros 50HQ, #82077 Thermo scientific). First, one volume of resin was washed with four volumes of 20 mM Tris-HCl pH 7.2. The use of vortex or centrifuge is not recommended as it can fragment the resin beads and block the column. Let decant 10 min before removing the supernatant. Resuspend again in four volumes of 20 mM Tris-HCl pH 7.2 and store at room temperature protecting from light. Poly-Prep^®^ Chromatography columns (Bio-Rad, 731-1550) were packed with 500 mg of resin for a protein amount range of 10-20 mg per sample. Resin was stacked by gravity for 10 min at room temperature before applying pressure using a hermetic cap and syringe to remove solution without drying out the resin. One volume of packed resin was equilibrated four times with two volumes of 20 mM Tris-HCl pH 7.2 under pressure. Peptide samples were diluted with one volume of 20 mM Tris-HCl pH 7.2 just before applying to the chromatography column. One volume of resin was washed four times with two volumes of cold 20 mM Tris-HCl pH 7.2, and then washed again with two volumes of 10 mM Tris-HCl, 20% ACN pH 7.2. Bound phosphopeptides were eluted from one volume resin with two volumes of 100 mM TEA (triethylamine) pH 11, and then with two volumes of 500 mM TEA pH 11. Residual liquid was collected from the resin, combined with the other eluates and the TEA evaporated using a speed-vacuum. Keep samples dry at −80°C until analysis.

### Immunoaffinity purification (IAP)

IAP columns containing one volume of protein A crosslinked with 1- and 3-pHis mAbs were equilibrated three times with one volume of NH_4_HCO_3_ pH 8.5 at RT without drying out the protein A.

Only if IAP is performed directly after digest in 50 mM NH_4_HCO_3_ pH 8,5 by trypsin, 100 µM of TLCK (Tosyl-L-lysyl-chloromethane hydrochloride, Trypsin inhibitor T7254 from Sigma) was added at RT 10 min before to pass the sample on column. A longer incubation is not recommended as the TLCK would be not stable over time in this condition (pH 8.5) decreasing its efficiency.

If samples were eluted from HAP/SAX, dried peptides are resuspended in 50 mM NH4HCO3 pH 8,5. Collecting the flow through, peptides were pass on column twice then washed five times with one volume of 50 mM NH4HCO3 pH 8,5. Finally, the phosphopeptides were eluted twice with one volume of 100 mM TEA pH 11. The phosphohistidine signal from elution can be controlled by peptide dot-blot. Elution sample were dried using speed-vacuum and store at −80°C until mass spectrometry analyses. The column can be washed immediately at least four times with one volume of PBS and stored with two volume of PBS at 4°C.

### LC-MS/MS

Samples are resuspended with cold buffer A (pH 3), on ice, just before desalting/loading under high pressure on Aqua-C18 5 µm resin (3 cm) within a 250 µm column (fused silica capillary with a Kasil frit). The same C18 resin (12 cm) is used for a 100 µm analytical column. Nano liquid chromatography tandem mass spectrometry (nano-LC-ESI MS/MS) was performed in positive ion mode using an Agilent 1200 G1311 Quat Pump coupled to an LTQ-Velos Orbitrap (CID35) or an Orbitrap Q-Exactive (HCD25). LC-gradient of 30 min, 90 min, 120 min and 180 min have been used with the buffer A and B corresponding to the following pH condition.

For pH 2 gradient: Buffer A (5% ACN, 95% H_2_O, 0.1% formic acid, pH 2), Buffer B (80% ACN, 20% H_2_O, 0.1% Formic Ac., pH 2).

For pH5 gradient: Buffer A (10mM Ammonium Bicarbonate, pH5), Buffer B (100% MetOH).

The 120-minute elution gradient had the following profile: 10% Buffer B at 5min, 40% Buffer B at 80 min, to 100% Buffer B at 100 min continuing to 110 min. The same profile was adapted for the different elution times. One cycle consisted of one full scan mass spectrum (m/z 300-1600) with a maximum injection time of 10 ms at 60,000 resolution followed by up to 20 data-dependent collision induced dissociation (CID) MS/MS spectra. Application of mass spectrometer scan functions and HPLC solvent gradients were controlled by the Xcalibur data system.

### In silico and data analysis

MS Raw Data were extracted using RawConverter. Phosphopeptide identification and localization of phosphates were done with MaxQuant (v1.6.2.6a) using Andromeda and IP2 (ProLuCID algorithm) with a Uniprot human database considering canonical and isoform reviewed sequences only ^2,42,43^. A reversed sequence library and contaminant list of peptides were integrated through MaxQuant software. Searches were performed using Trypsin/P digest with 3 max missed cleavages due to potential phosphorylation of both tryptic cleavage sites, Lys and Arg. Variable modifications: Oxidation (M); Acetyl (Protein N-term); Phospho (STY)(HKR-98) “Not C-ter”, meaning the use of default parameters for S, T, Y phosphoresidues and the additional neutral loss of 97.9768950947 (HPO_3_ + H_2_O) for H, K, R residues excluding all C-terminal phosphorylations. Fixed modifications: Carbamidomethyl (C). A total of 5 modifications maximum were considered per peptides.

For synthetic peptides analysis (Supplementary Fig. 2), decoy enzymes were created to consider in-silico the correct sequences from the Uniprot human database and obtain probability of localization data: Histone H4 (cleaves G.G and L.R pairs), PGAM1 (cleaves A.W and L.V pairs), NME1/2 (cleaves G.R and S.V pairs). Amidated (C-ter) was integrated as a modification for the correct mass assignment of these synthetic peptides and no miscleavages were allowed.

First search peptide tolerance at 20 ppm and main search at 4.5 ppm. Andromeda parameters were used by default with a minimal peptide length of 7 residues and maximum peptide mass of 4,600 Da. False discovery rate (FDR) fixed at 0.01 and minimum ion score for modified peptides at 40. ITMS MS/MS match tolerance was of 0.5 Da and FTMS MS/MS match tolerance was of 20 ppm.

The consideration of all SONAtes residues S, T, Y, H, K, R, D, E, C (Supplementary Fig. 4B) required the use of a new variable modification Phospho(STY)(HKRDEC-98) Not C-ter in MaxQuant, corresponding to the previous Phospho(STY)(HKR-98) Not C-ter modification with the addition of D, E, C residues, with a similar neutral loss of 97.9768950947 (HPO_3_ + H_2_O) within the specificity for the configuration of Andromeda.

Complement analysis with histidine, lysine and arginine immonium ions, not shown, considered the additional diagnostic peaks for pHis (neutral mass: 190.0376 m/z), for pLys and pLys less ammonia (neutral mass respectively: 181.0737 m/z and 164.0471 m/z) and for pArg (neutral mass: 209.0798 m/z).

All MaxQuant outputs txt files were processed with Perseus software (v1.6.2.1) to list and select significant localization of phosphate. Finally, data were exported on Excel sheet for graphic interpretation. The final list of 425 phosphoramidate residues were analyzed on Weblogo (v3.6.0) to observe a potential motif and a GO molecular function analysis was made using Fisher test type with both FDR and Bonferroni correction (Supplementary Table 3).

#### Protein fractionation

PC-3 cells were used to test the compatibility of a neutral fractionation method. Protein concentration of 2.7 mg/mL was defined by BioRad protein assay. 500 µg protein at 1 mg/mL were prepared with Buffer A (Phosphoprotein Kit, Clontech, Cat#635626): 185 µL PC-3 lysate (pH 8.8), 65 µL Buffer A (pH 6), 250 µL Water (HPLC grade). The fast fractionation solution (FF) used corresponded to 90% MeOH (HPLC grade), 10% Water (HPLC grade) and each fraction were collected as following:

Fraction #1: 7.5 µL FF + 500ul PC-3 sample

Fraction #2: 8.0 µL FF + Supernatant from Fraction 1

Fraction #3: 16.5 µL FF + Supernatant from Fraction 2

Fraction #4: 25.0 µL FF + Supernatant from Fraction 3

Fraction #5: 32.5 µL FF + Supernatant from Fraction 4

Fraction #6: 40.0 µL FF + Supernatant from Fraction 5

Fraction #7: 85.0 µL FF + Supernatant from Fraction 6

Fraction #8: 130.0 µL FF + Supernatant from Fraction 7

Fraction #9: 425.0 µL FF + Supernatant from Fraction 8

Fraction #10-1: 850.0 µL FF + Supernatant from Fraction 9(50%)

Fraction #10-2: 850.0 µL FF + Supernatant from Fraction 9(50%)

Each fraction was centrifuged in the cold room for 10 min at maximum rpm and the peptides were identified on a Fusion Orbitrap with CID35 fragmentation and a 140 min LC-gradient. Spectral counts and peptide counts were extracted from IP2 platform analysis.

#### Raw Data file

A list of experiment, samples and run for each sample is provided and assign for every Raw file (Supplementary Table 4).

## Acknowledgements

The authors are grateful for financial support from the George E. Hewitt Foundation for Medical Research for providing K. Adam with a postdoctoral fellowship, T. Hunter is a Frank and Else Schilling American Cancer Society Professor, and holds the Renato Dulbecco Chair in Cancer Research. NIH support for these studies was from the NCI R01 CA080100, CA082683, CA194584 to T.H.; NIGMS GM103533 and NCRR 5P41RR011823 to J.R.Y. III; and CCSG support for the Mass Spectrometry Core Service was from NCI CA014195.

## Author contributions

K. Adam conceived, designed and performed all experiments of the study and wrote the paper, S. Fuhs generated and provided the hybridoma clones used to make anti-1/3 pHis monoclonal antibodies, A. Aslanian contributed to early stage development for direct injection of synthetic peptides for MS analysis, J Meisenhelder made the synthetic peptides and gave technical advice, J. Diedrich and J. Moresco performed some of the MS runs on the Q-Exactive and Fusion Orbitrap instruments with HCD fragmentation and the methanol gradient fractionation experiment, J. La Clair provided material assistance and expertise for phosphoramidate synthesis, J.R. Yates III provided access to a LTQ-Velos Orbitrap mass spectrometer and insights, T. Hunter provided laboratory equipment, scientific guidance and edited the paper.

### Competing interests

The authors declare no competing interests.

---

**Sup. Fig. 1:**
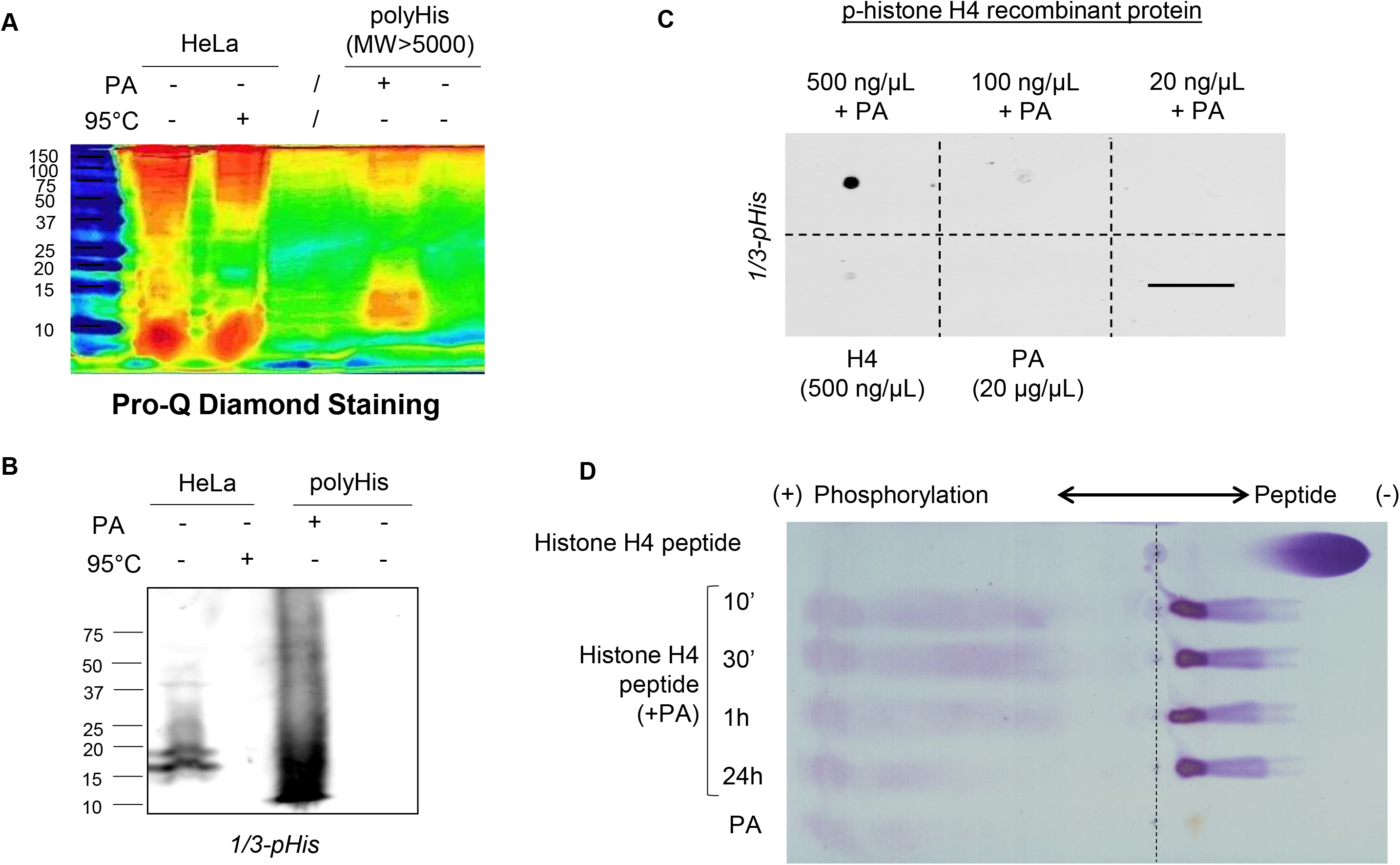
Validation of N-phosphate chemical phosphorylation with phosphoramidate (PA). **A**. Pro-Q Diamond in-gel staining reveals phosphorylations from HeLa cells and chemical phosphorylation on polyhistidine peptides. Signal is decreased or lost after boiling samples with the thermosensitive pHis. **B**. A mix of 1/3-pHis antibodies were used to reveal polyhistidine chemically phosphorylated by immunobloting. **C**. A dot blot of histone H4 recombinant protein after chemical phosphorylation with phosphoramidate (PA) reveals histidine phosphorylation using 1/3-pHis, or specific phospho-histone H4 (H18) antibodies (data not shown). **D**. Thin-Layer Electrophoresis (TLE) at pH 8.9 of histone H4 (GAKRHRKVL) peptide, confirm non-conventional phosphorylation.

**Sup. Fig. 2:**
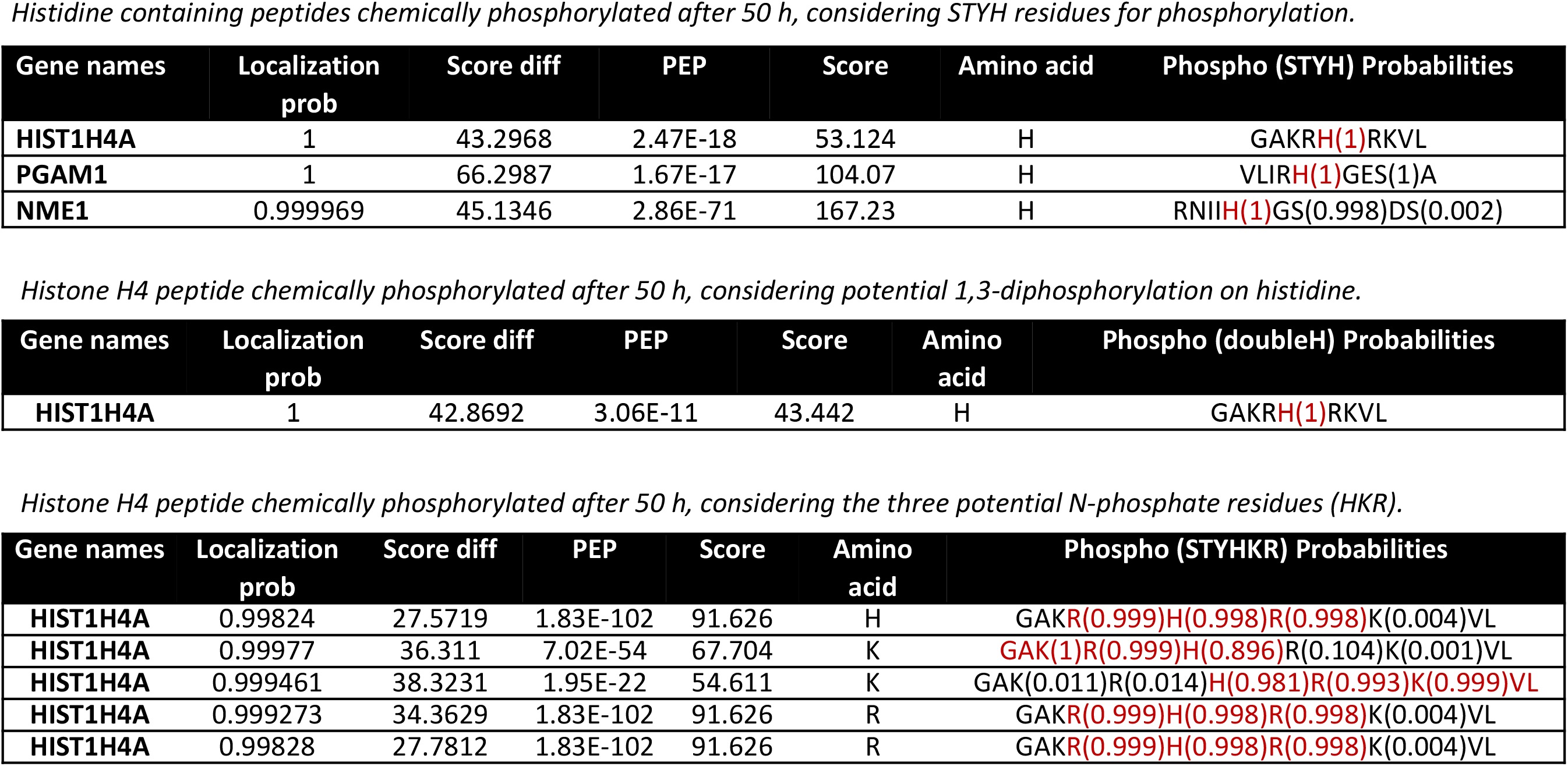
MaxQuant identification of chemically phosphorylated synthetic peptides after 50 h reaction. Top table corresponds to the search for p-histone H4, p-PGAM1 and p-NME1 considering only conventional STY phosphoresidues and phosphorylated histidine (+98 Da neutral loss). Second table corresponds to the search of potential 1,3-diphospho-His on histone H4 peptide. The third table integrates all the conventional O-phosphate (STY) and the three N-phosphate bonds (HKR, +98 Da neutral loss).

**Sup. Fig. 3:**
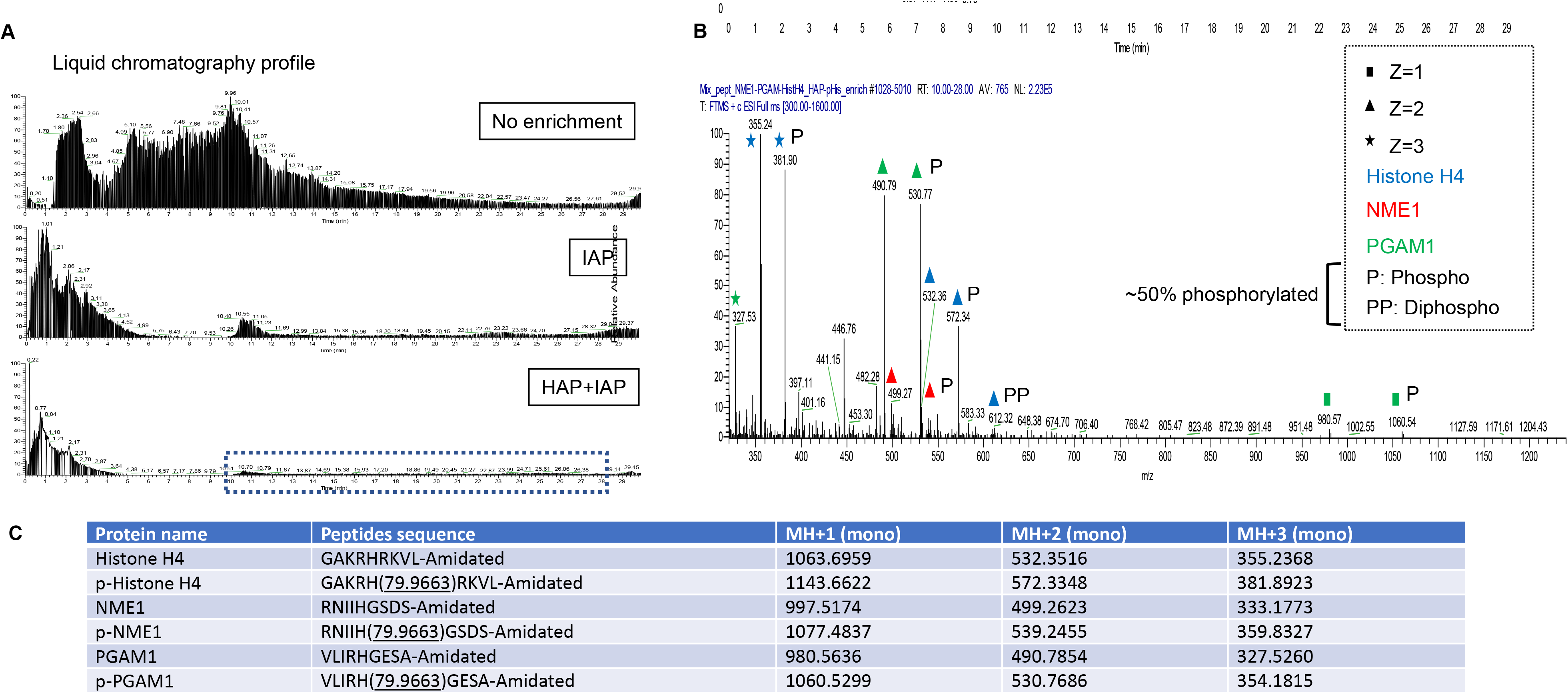
LC-MS/MS analysis of chemically phosphorylated synthetic peptides after alkaline pHis enrichment. **A**. Three liquid-chromatography profiles represent the pool of synthetic peptides (NME1, PGAM1, histone H4) chemically phosphorylated with no enrichment, after 1/3-pHis ImmunoAffinity Purification (IAP) alone, or successive HAP+IAP, from top to bottom respectively. HAP for Hydroxyapatite. **B**. MS profile from combined HAP+IAP enrichment (retention time window: 10-28 min). Peptide sequences are indicated through color blue (histone H4), red (NME1) and green (PGAM1), and the charge state (z) is symbolized by a square (+1), triangle (+2) or a star (+3). Peaks corresponding to the phosphorylation forms are manually annotated with P or PP. **C**. Theoretical monoisotopic mass/charge (m/z) ratio for each non-phosphorylated and phosphorylated peptide sequences used for manual annotations. (using MS-product from UCSF - Protein Prospector V5.20.0 - Baker, P.R. and Clauser, R. http://prospector.ucsf.edu).

**Sup. Fig. 4:**
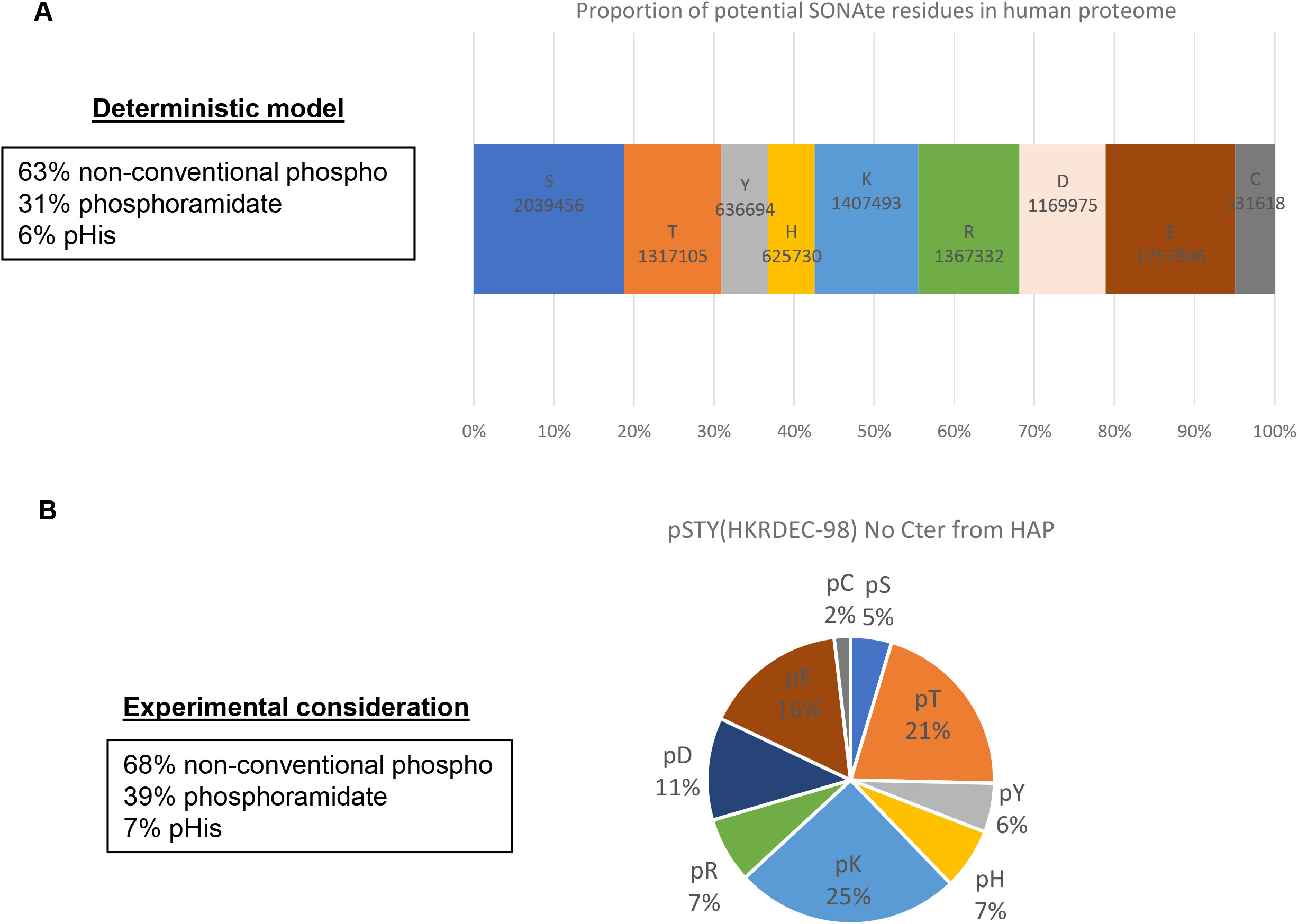
**A**. The deterministic model represented here considers the total amount for each of the SONAte residues (STYHKRDEC) in human proteome as initial condition. The same probability of phosphorylation for each one of these residues, without random influence of protein conformation, sequence motif, various stability or recovery, were used as determined parameter. **B**. Experimental consideration of every SONAte residues for identification from from HeLa cells after global phosphopeptide enrichment without any specificity in non-acidic condition.

**Sup. Fig. 5:**
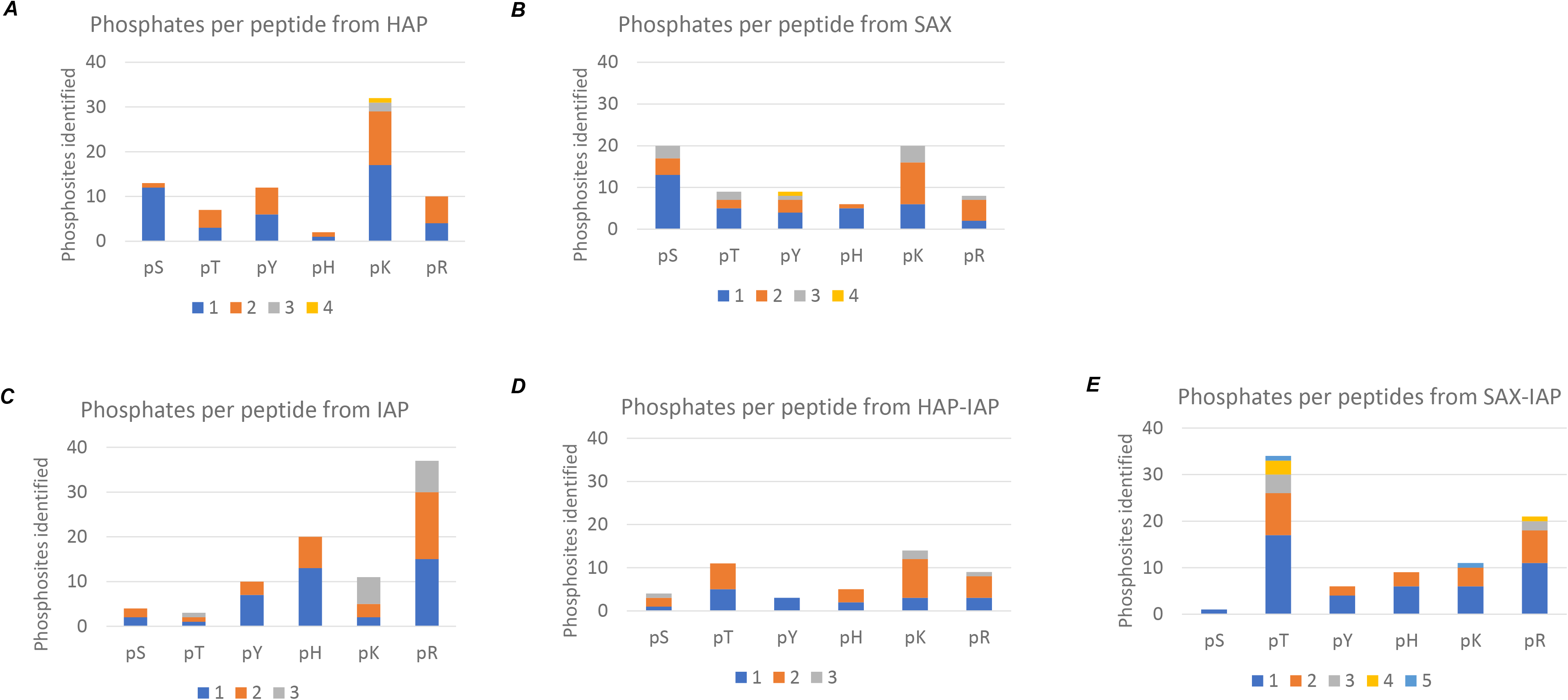
Numbers of each phosphoresidue localized with a high probability within a peptides with 1 to 5 phosphates. **A. and B**. Global phosphopeptide enrichment from HAP or SAX chromatography. **C**. Unique IAP enrichment using 6 combined pHis mAbs (1-pHis clones: SC1-1, SC50-3, SC77-11; 3-pHis clones: SC39-6, SC44-1, SC56-2). **D. and E**. Combination of global phosphopeptides enrichment followed by 1/3-pHis immunoaffinity purification

**Sup. Fig. 6:**
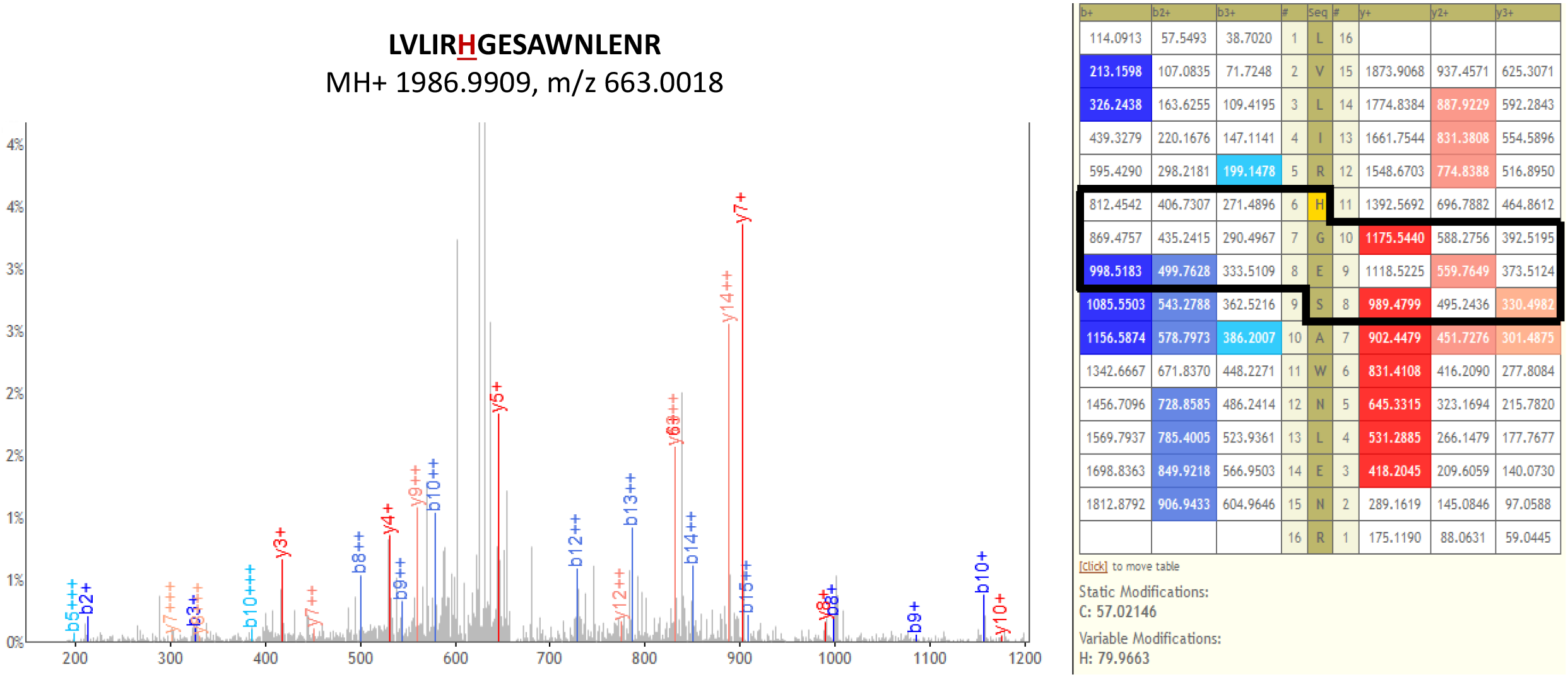
Analyses from integrated proteomic pipeline IP2 confirmed the position and revealed specific peaks (b/y ions) for pHis compare to the closest Ser. 6 peaks MS/MS (b/y ions) detected with charges +1, +2 and +3 are specific for pHis position versus a potential pSer site.

**Sup. Fig. 7:**
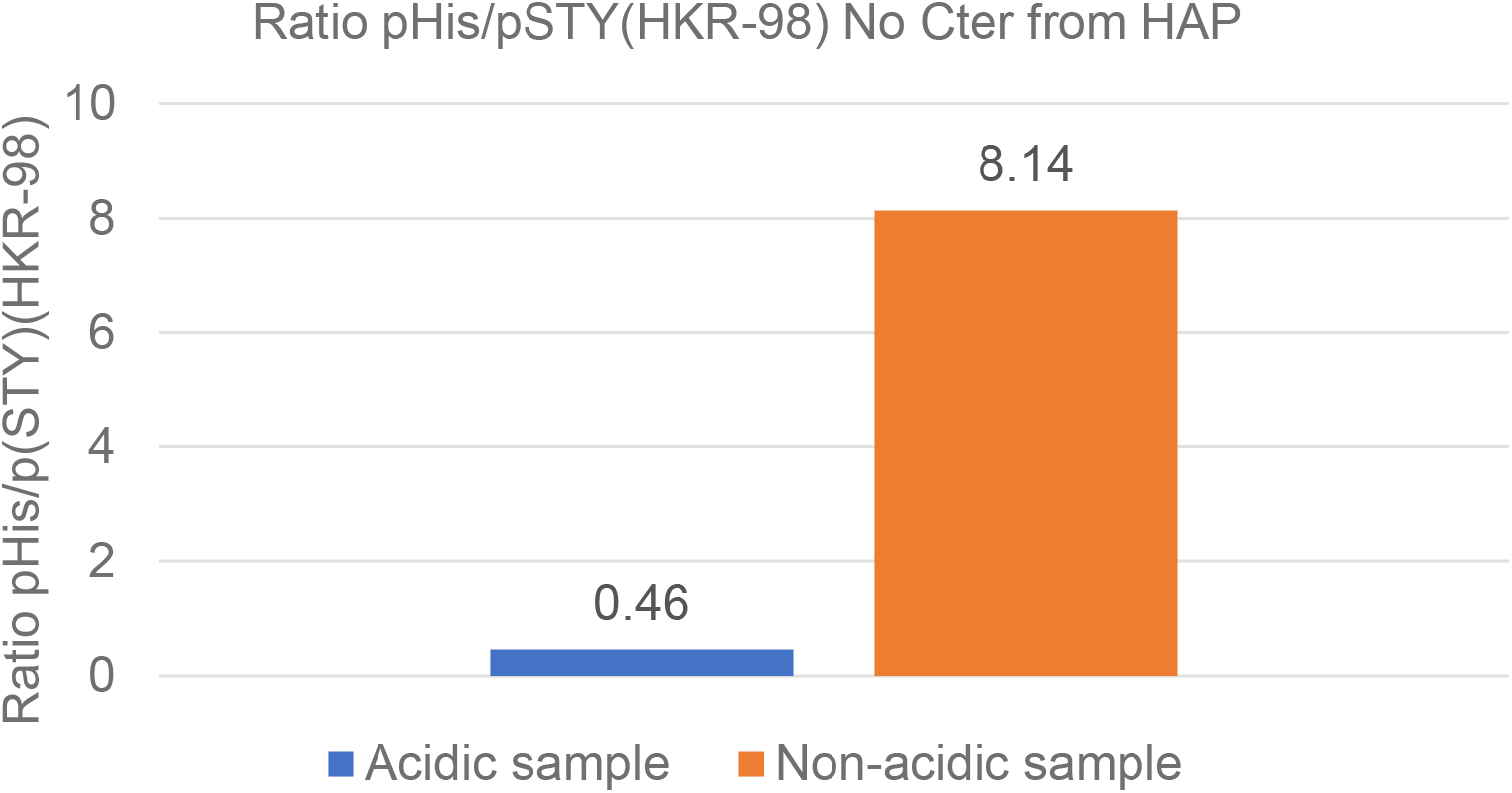
Comparative analysis of acidic versus non-acidic sample for HAP global phosphopeptide enrichment under non-acidic conditions. After identification of S, T, Y, H, K, R phosphorylation using MaxQuant configuration, the ratio of pHis sites and total phosphorylation sites was defined.

**Sup. Fig. 8:**
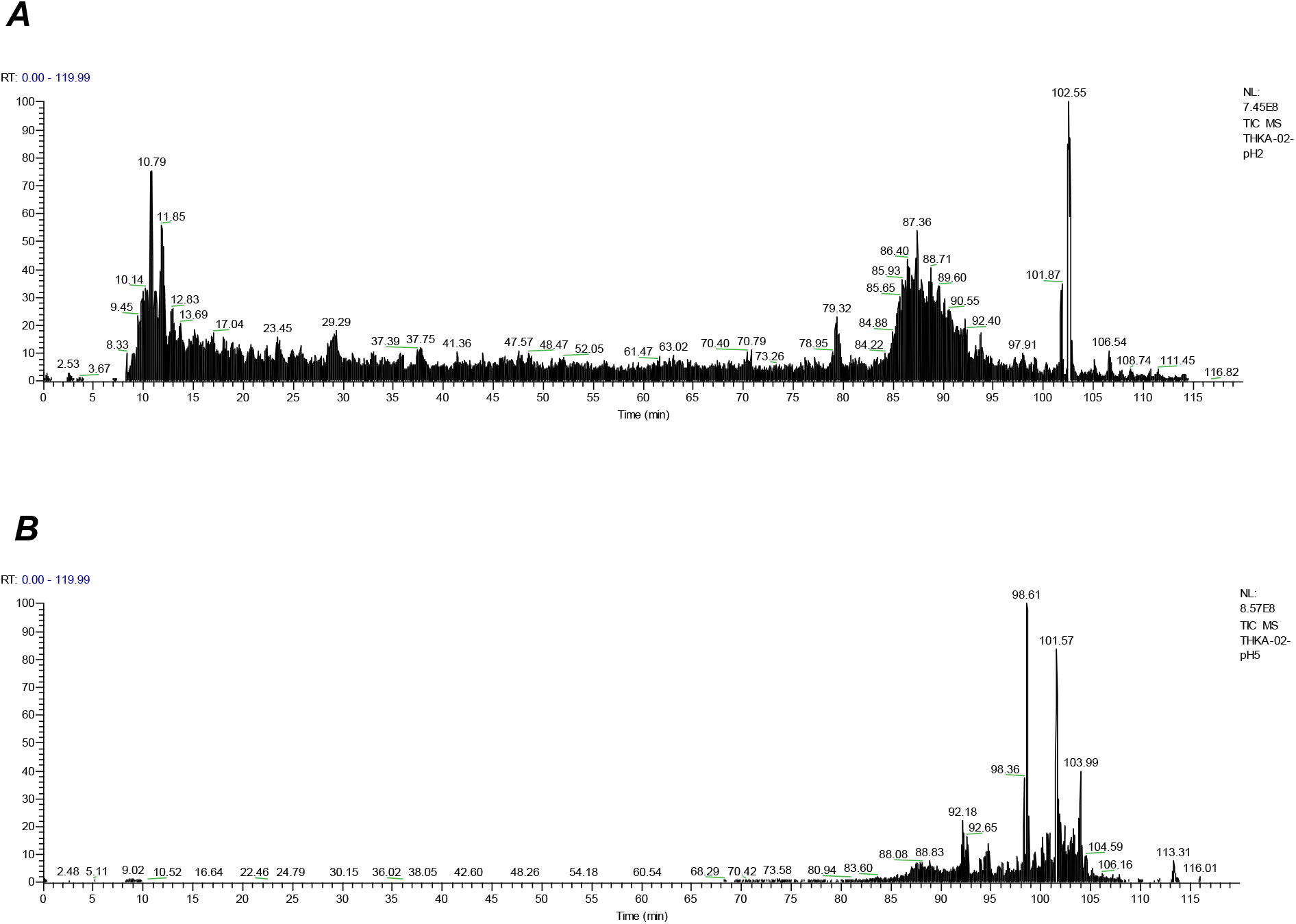
ImpactofhigherpHonLC-MSprofile.Thesame3-pHisIAPsample(THKA-02)wereusedfortwocomparativeionizationconditionswithnESIbufferandchromatographyatpH2(**A**)and pH5(**B**).

**Sup. Fig. 9:**
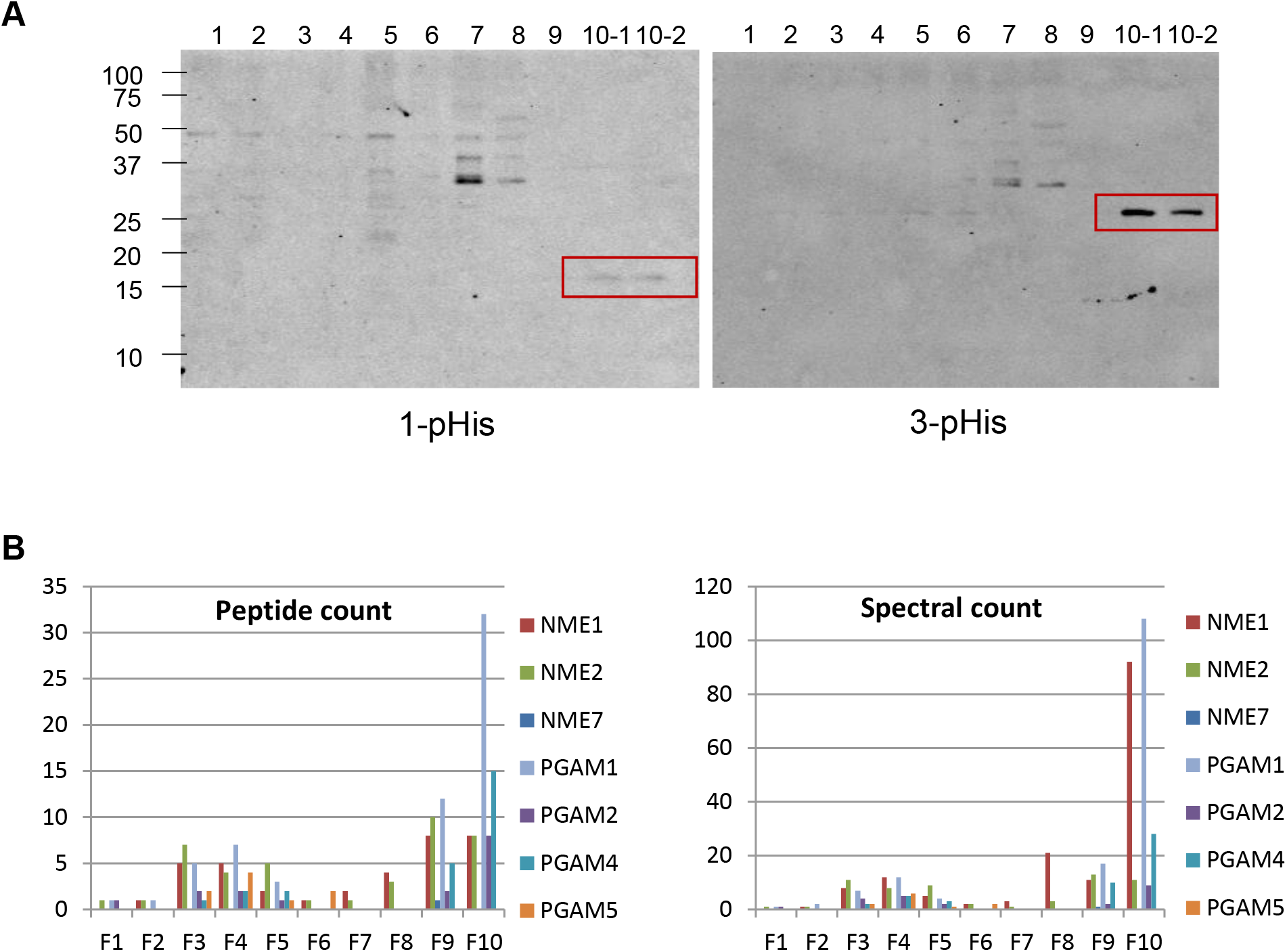
Compatibility and efficiency of fast protein fractionation under neutral conditions (90% Methanol) from human cancer cells PC-3 lysates. **A**. Immunoblot showing the 1-pHis and 3-pHis containing proteins from each fraction on the left and right panel, respectively. The red rectangle highlights the corresponding molecular weight of known 1-pHis NME1/2 and 3-pHis PGAM1. **B**. After MS identification of NME and PGAM family members, the respective peptide count and spectral count were defined for each fraction (F1-F10).

